# CNTN4 modulates neural elongation through interplay with APP

**DOI:** 10.1101/2023.08.25.554833

**Authors:** Rosemary A. Bamford, Amila Zuko, Jan J. Sprengers, Harm Post, Renske L. R. E. Taggenbrock, Annika Mehr, Owen J. R. Jones, Aurimas Kudzinskas, Josan Gandawijaya, Madeline Eve, Ulrike C. Müller, Martien J. Kas, J. Peter H. Burbach, Asami Oguro-Ando

## Abstract

The neuronal cell adhesion molecule contactin-4 (CNTN4) has been genetically linked to autism spectrum disorders (ASD) and other psychiatric disorders. The *Cntn4*-deficient mouse model has previously shown that CNTN4 has important roles in axon guidance and synaptic plasticity in the hippocampus. However, the pathogenesis and functional role of CNTN4 in the cortex have not yet been investigated.

Using Nissl staining, immunohistochemistry and Golgi staining the motor cortex of *Cntn4^-/-^* mice was analysed for abnormalities. Interacting partners of CNTN4 were identified by immunoprecipitation and mass spectrometry. Further analysis of the interaction between CNTN4 and APP utilised knockout human cells generated via CRISPR-Cas9 gene editing.

Our study newly identified reduced cortical thickness in the motor cortex of *Cntn4^-/-^* mice, but cortical cell migration and differentiation were unaffected. Significant morphological changes were observed in neurons in the M1 region of the motor cortex, indicating that CNTN4 is also involved in the morphology and spine density of neurons in the motor cortex. Furthermore, mass spectrometry analysis identified an interaction partner for CNTN4, and we confirmed an interaction between CNTN4 and APP. Knockout human cells of CNTN4 and/or APP revealed a relationship between CNTN4 and APP.

This study demonstrates that CNTN4 contributes to cortical development, and that its binding and interplay with APP controls neural elongation. This is an important finding for understanding the function of APP, a target protein for Alzheimer’s disease. The binding between Cntn4 and APP, which is involved in neurodevelopment, is essential for healthy nerve outgrowth.

## Introduction

Neurodevelopmental disorders such as Autism spectrum disorder (ASD) are highly heterogeneous in terms of genetics, behaviour, and pathology (1–3). Autism spectrum disorder (ASD) is characterised by core behavioural domains such as impaired social communication and interactions, repetitive behaviour and restricted interests (1,2). A clinical diagnosis of ASD is based primarily upon behaviour, however ASD encompasses a range of syndromes and severity (3). Over a 1000 genes are implicated in ASD according to the Simons Foundation Autism Research Initiative (4) and this list is still growing. However, the mechanisms of how current ASD associated genes (4) influence neurodevelopment and are involved in ASD are not fully understood. It has been proposed that there are four converging molecular processes important to these genes: neuronal communication (5), neuronal cell adhesion (6), excitation and inhibition balance (7,8), and regulation of post-synaptic translation (9). Neuronal cell adhesion gene networks, in particular those that are acting in the synaptic complex (10,11), are evidenced to be involved in ASD pathogenesis, and well-studied genes include neurexin-1 (*NRXN1*) (12), neuroligin-1 (*NLGN1*) (13,14), neuroligin-4 (*NLGN4*) (15) and contactin associated protein-2 (*CNTNAP2*) (16–19). Cell adhesion molecules are integral to neuronal migration, axon guidance and neuron-glial cell interactions; processes important for cortical development and often disturbed in ASD (20–22). ASD has been associated with abnormalities in the cerebral cortex and subsequently affected cognitive functions (i.e. communication, interaction and learning) (23) and multiple pathologies in cortical areas (24–26).

Contactins are a protein family belonging to a specific subclass of the immunoglobulin cell adhesion molecule superfamily (IgCAM). These proteins share 40-60% homology at the amino acid sequence level (27). Contactins have been previously associated with neurodevelopmental disorders and constitute an interesting group of proteins to investigate in relation to ASD aetiology (28–30). Copy number variants (CNVs) have been identified in cell adhesion molecules in ASD patients, which further suggests a role for this group of proteins in ASD development (31–34). Among them, Contactin 4 (CNTN4) and Contactin 6 (CNTN6), have been reported as candidate genes for chromosome 3 disorders (6), particularly CNTN4, which has been previously linked to genetic, behaviour and pathological studies of ASD (35–38).

Cntn4 is mainly expressed in the cerebral cortex layers II-V in addition to the olfactory bulb, thalamus and hippocampus (38) as well as expressed on cortical pyramidal and interneurons (39). The domains of human CNTN4 are conserved in mice (40), therefore mouse models can be utilised to reveal the role of CNTN4 in normal and abnormal development. Kaneko-Goto *et al.* generated *Cntn4*-deficient mice and observed the importance of CNTN4 as an axon guidance molecule in the olfactory bulb (41), with further evidence in the optic system (42). Therefore, CNTN4 acts as one of the axon guidance molecules crucial for the proper formation and development of the olfactory and optic systems (41,42). More recently, Molenhuis *et al.* used a developmental behaviour battery of Cntn4-deficient mice to reveal subtle non-disease-specific changes in sensory behavioural responses and cognitive abilities (43), while Oguro-Ando *et al.* showed that CNTN4 is associated with morphological and synaptic plasticity changes in hippocampal CA1 neurons and has important functions in fear memory in their experiments using *Cntn4*-deficient mice (44). However, since CNTN4 lacks transmembrane and intracellular domains, it is closely dependent on *cis* or *trans* interactions with membrane-spanning proteins (27). The full extent of the molecular network with which CNTN4 interacts, as well as the role of CNTN4 in cortical and neuronal development, is still not clear.

The aim of this study was to identify novel functions of CNTN4 in the cortex at protein, cellular and anatomical levels using *Cntn4^-/-^* mice. This provides information on the role CNTN4 has in neurodevelopmental processes. We investigated the phenotype in the cortex of the *Cntn4^-/-^* mice and found that the disruption of neuroanatomical organisation caused as a consequence of CNTN4 deficiency was confirmed by structural and morphological changes in the cortex. We then examined the binding partners of CNTN4 and the impact of their interactions on the expression levels and morphology of human cell lines, suggesting that CNTN4 plays an important role in the motor cortex, which may lead to further insights into the aetiology of ASD and neurodevelopmental mechanisms.

## Results

### Abnormal cortical thickness observed in Cntn4-/-mice

In the cerebral cortex, the cortical layer thickness is involved in migration and may be an indicator of neurodevelopment abnormalities. Nissl staining and microscopic imaging of brain sections were carried out in order to measure the cortical layer thickness of *Cntn4^-/-^*, *Cntn4^+/-^* and *Cntn4^+/+^* mice, respectively (Figure 1A). The thickness of the upper layers (I-IV), lower (V-VI) layers and the total thickness were measured for the motor cortex, somatosensory cortex and visual cortex. Quantitative results showed that the cortical thickness of all layers for the somatosensory and visual cortices did not differ between the genotypes (Figure S1, p > 0.05, one-way ANOVA). However, the thickness of the upper layer of the motor cortex was significantly reduced in the Cntn4^-/-^ mice (Figure 1B, p = 0.0188, one-way ANOVA). Therefore, Cntn4 deficiency leads to cortical thinning of the primary motor cortex, indicating abnormal gross cortical development.

**Figure 1:**
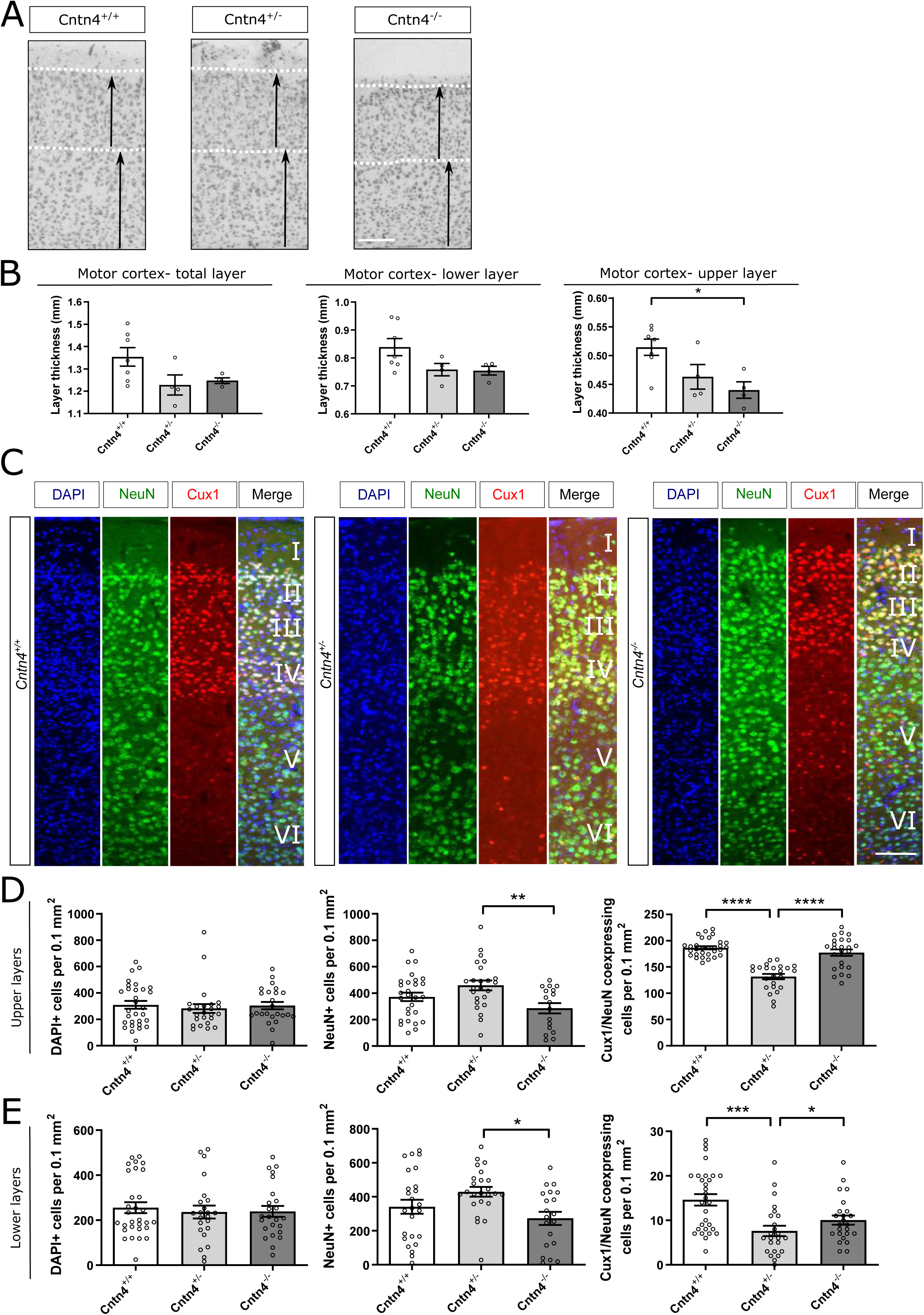
Cortical thickness and upper layer pyramidal neurons in *Cntn4^-/-^* mice. (A) Nissl-stained sections of the motor cortex of adult *Cntn4^+/+^*, *Cntn4^+/-^* and *Cntn4^-/-^* mice. Arrows indicate upper and lower layer thickness. The scale bar indicates 250 µm. (B) Quantitative analysis of the represented cortical layer thickness (upper I-IV, lower V-VI and total) demonstrate a significant difference in motor cortex thickness in all layers between *Cntn4^+/+^* and *Cntn4^-/-^* mice. Analysis was performed, in each area, on at least three slices. Data are presented as mean ± S. E. M, *Cntn4^+/+^* (n=7), *Cntn4^+/-^* (n=4) and *Cntn4^-/-^* mice (n=4), p = 0.0188 (upper layer, *Cntn4^+/+^* versus *Cntn4^-/-^*). (C) Images of Cux1 expression (red) in layers II-IV of the motor cortex together with the NeuN (green) in adult *Cntn4^+/+^*, *Cntn4^+/-^* and *Cntn4^-/-^* mice. DAPI is in blue. The scale bar represents 50 µm. (D) Quantitative analysis of the upper layers (layers II-IV) did not show differences in total cell number (p > 0.05, one-way ANOVA), but did show significant differences in total neuron number and Cux1+ neurons in the *Cntn4^+/-^* mice (p = 0.0058 and p < 0.0001, respectively, one-way ANOVA). (E) Quantitative analysis of the lower layers (layers V-VI) did not show differences in total cell number (p > 0.05, one-way ANOVA), but did show significant differences in total neuron number (p = 0.013, one-way ANOVA) and Cux1+ neurons (p = 0.0201, *Cntn4^+/+^* vs *Cntn4^-/-^*; p = 0.0002, *Cntn4^+/+^* vs *Cntn4^+/-^*, one-way ANOVA) in the *Cntn4^+/-^* mice. Analysis was performed, in each area, on at least six slices in *Cntn4^+/+^* (n=5), *Cntn4^+/-^* (n=4) and *Cntn4^-/-^* mice (n=4) using unpaired Student’s *t* test. Data are presented as mean ± S. E. M.

### Tissue architecture and cell differentiation influenced by loss of *Cntn4*

Next, we focussed on the laminar positioning and numbers of specific cortical pyramidal neurons in the motor cortex of *Cntn4^+/+^*, *Cntn4^+/-^* and *Cntn4^-/-^* mice. The location and density of upper- and lower-layer pyramidal neurons were investigated by visualisation of neurons destined for layers II-IV (red, Figure 1C) in adult mice using Cux1 as a marker. Cell counting revealed a significant decrease in the total number of neurons, which are NeuN+ (green, Figure 1C, p = 0.0058, one-way ANOVA) in the upper layers (layers II-IV), between *Cntn4^+/-^* and *Cntn4^-/-^* mice (Figure 1D). However, there was no significant difference in total number of cells, which are DAPI+ (blue, Figure 1C, p > 0.05, one-way ANOVA) in the upper layers between genotypes. There was a significant increase in Cux1/NeuN+ neurons in the upper layers for *Cntn4^+/+^* and *Cntn4^-/-^* mice with respect to *Cntn4^+/-^* mice (Figure 1D, p < 0.0001, one-way ANOVA). Measurements revealed no statistical difference in the total numbers of cells, neurons (Figure 1E, p > 0.05, one-way ANOVA). There was a significant decrease in the number of Cux1+ displaced neurons in the lower layers (layers V-VI) of *Cntn4^-/-^* mice (Figure 1E, p = 0.0201, *Cntn4^+/+^* vs *Cntn4^-/-^*; p = 0.0002, *Cntn4^+/+^* vs *Cntn4^+/-^*, one-way ANOVA). Similar to the upper layers, a significant decrease was observed in the total number of neurons which are NeuN+ (green, Figure 1C) in the lower layers, between *Cntn4^+/-^* and *Cntn4^-/-^* mice (Figure 1E, p = 0.013, one-way ANOVA). Overall, neuronal density and cortical migration differed in the M1 region but does not account for the cortical thinning observed in the Nissl staining. These results indicate that CNTN4 is required for normal development of pyramidal neurons in the upper layers of the motor cortex.

### *Cntn4* deficiency affects dendrite length and complexity

In parallel to investigating the density and migration of CNTN4 deficient cortical pyramidal neurons, the morphology of Golgi-stained pyramidal neurons in the motor cortex layers II-III were analysed for dendrite length, branching and complexity in the *Cntn4^+/+^, Cntn4^+/-^* and *Cntn4^-/-^* mice (Figure 2A-B). The longest dendrite length was significantly decreased in the apical dendrites of *Cntn4^-/-^* mice compared to the *Cntn4^+/-^* mice (p = 0.0245, one-way ANOVA) (Figure 2C).

**Figure 2:**
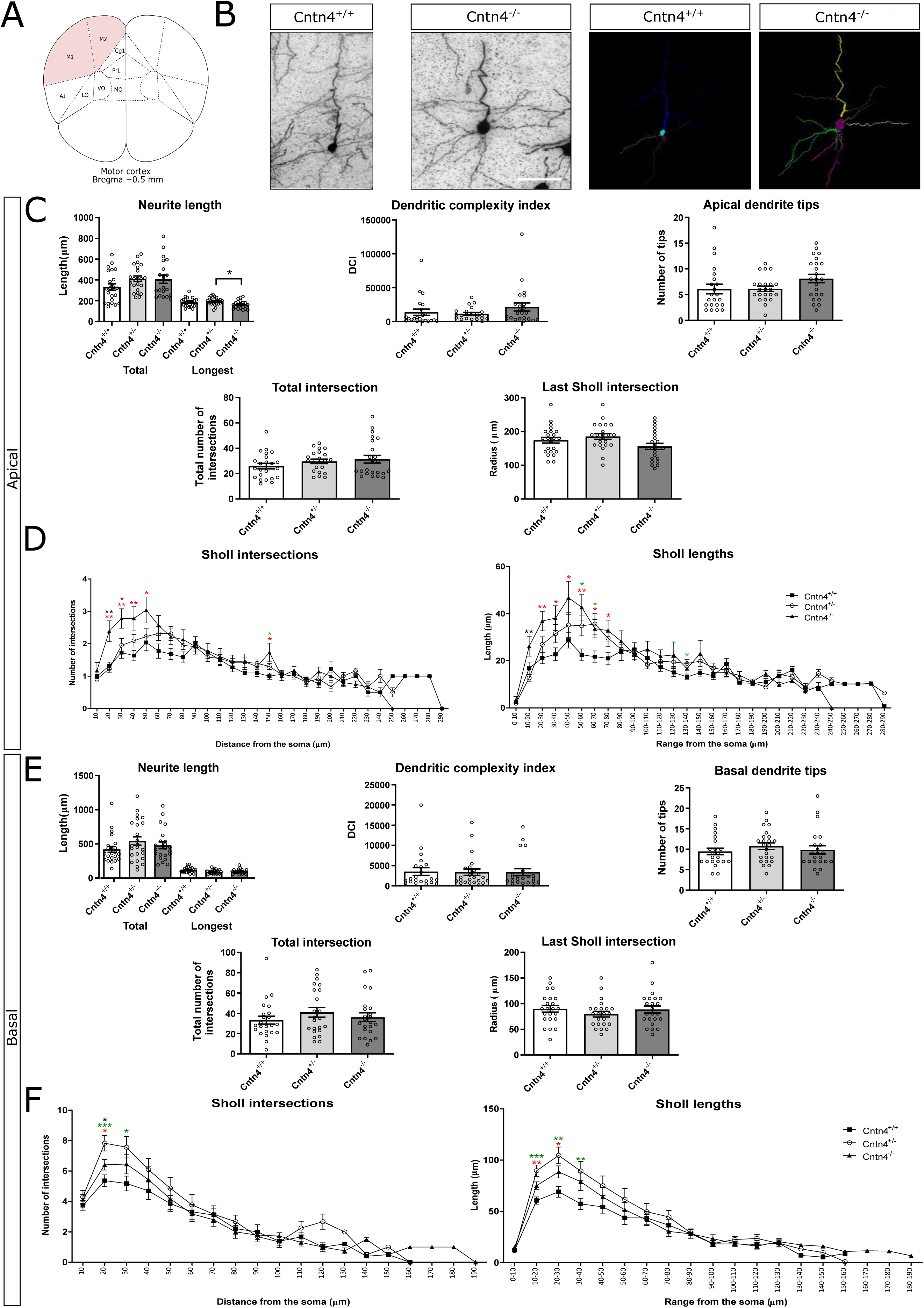
Neuron morphology analysis results for *Cntn4^+/+^*, *Cntn4^+/-^* and *Cntn4^-/-^* mice primary motor cortex, layers II-III. A) Schematic representation of the motor cortex with labelled Bregma anterior-posterior. Adapted from Paxinos and Franklin, 2001. B) Golgi staining in *Cntn4^+/+^* and *Cntn4^-/-^* mice motor cortex and trace outlines of example pyramidal neurons. The scale bar represents 40 µm. The arrowheads show differences. C) & E) Quantitative morphological results for the apical and basal dendrites respectively. D) & F) Sholl plots indicate the distribution of respective apical and basal dendritic intersections and length at increasing distance from the centre of the cell body. Quantitative analysis was performed, in each area, on at least three slices in *Cntn4^+/+^*, *Cntn4^+/-^* and *Cntn4^-/-^* mice (n=5 per genotype) using one-way ANOVA. Data are presented as mean ± S. E. M, cell number n=23 (*Cntn4^+/+^*), n=22 (*Cntn4^+/-^*), n=23 (*Cntn4^-/-^*) (p = 0.0245, *Cntn4^+/-^* vs *Cntn4^-/-^*, longest apical neurite length). Red asterix corresponds to *Cntn4^+/+^* versus *Cntn4^-/-^*; green asterix corresponds to *Cntn4^+/+^* versus *Cntn4^+/-^* and black asterix corresponds to *Cntn4^-/-^* versus *Cntn4^+/-^*, * p < 0.05.

The dendritic complexity index (DCI) was calculated in order to compare the complexity of the branching dendrites and length between pyramidal neurons. The more complex the branching structure, or longer the dendrites, the higher the index is. For the apical and basal dendrites an equal complexity was observed across all genotypes (Figure 2C, E, p > 0.05, one-way ANOVA). The number of apical and basal dendrite tips, and total intersections, was not significantly different between genotypes (Figure 2C, E, p > 0.05, one-way ANOVA).

Sholl plots indicate the distribution of dendritic intersections and length at increasing distance from the centre of the cell body (45). The last Sholl apical and basal intersections were not significantly different between genotypes (Figure 2C, E, p > 0.05, one-way ANOVA). It was observed that the *Cntn4^-/-^* mice had a significantly higher number of Sholl apical intersections in the range of 50 µm from the soma, compared to *Cntn4^+/+^* mice. The *Cntn4^-/-^* mice had significantly longer Sholl apical lengths in the range of 80 µm from the soma, compared to the *Cntn4^+/+^* mice (Figure 2D, p < 0.05, one-way ANOVA). It was observed that the *Cntn4^-/-^* mice had a significantly higher number of Sholl basal intersections within 20 µm from the soma, compared to *Cntn4^+/+^* mice. The *Cntn4^-/-^* mice had significantly longer Sholl basal lengths within 30 µm from the soma, compared to the *Cntn4^+/+^* mice (Figure 2F, p < 0.05, one-way ANOVA). These data indicate that CNTN4 deficiency causes abnormal apical and basal neurite growth in cortical pyramidal neurons.

### Spine number and maturity are changed in *Cntn4^-/-^* mice

To further investigate the role of CNTN4 in the cell function and fate of pyramidal neurons in the motor cortex, we employed Golgi staining to assess spine numbers and their maturity status (Figure 3A). Loss of CNTN4 significantly increased the total number of spines 50 µm from the cell soma, compared to *Cntn4^+/+^* mice (p=0.0057) (Figure 3B). The maturity of spines also differed for the *Cntn4*-deficient mice; there were significantly fewer immature spines (A type) and more abnormal spines (F/G types) (Figure 3C-E). These differences were more pronounced in the first 25 µm than in the latter 25 µm. Expression of the other maturity morphologies was comparable between the genotypes. Since the total number of spines increases overall, this implies that loss of *Cntn4* expression leads to an increase of abnormal spines at the expense of immature spines which decrease in number.

**Figure 3:**
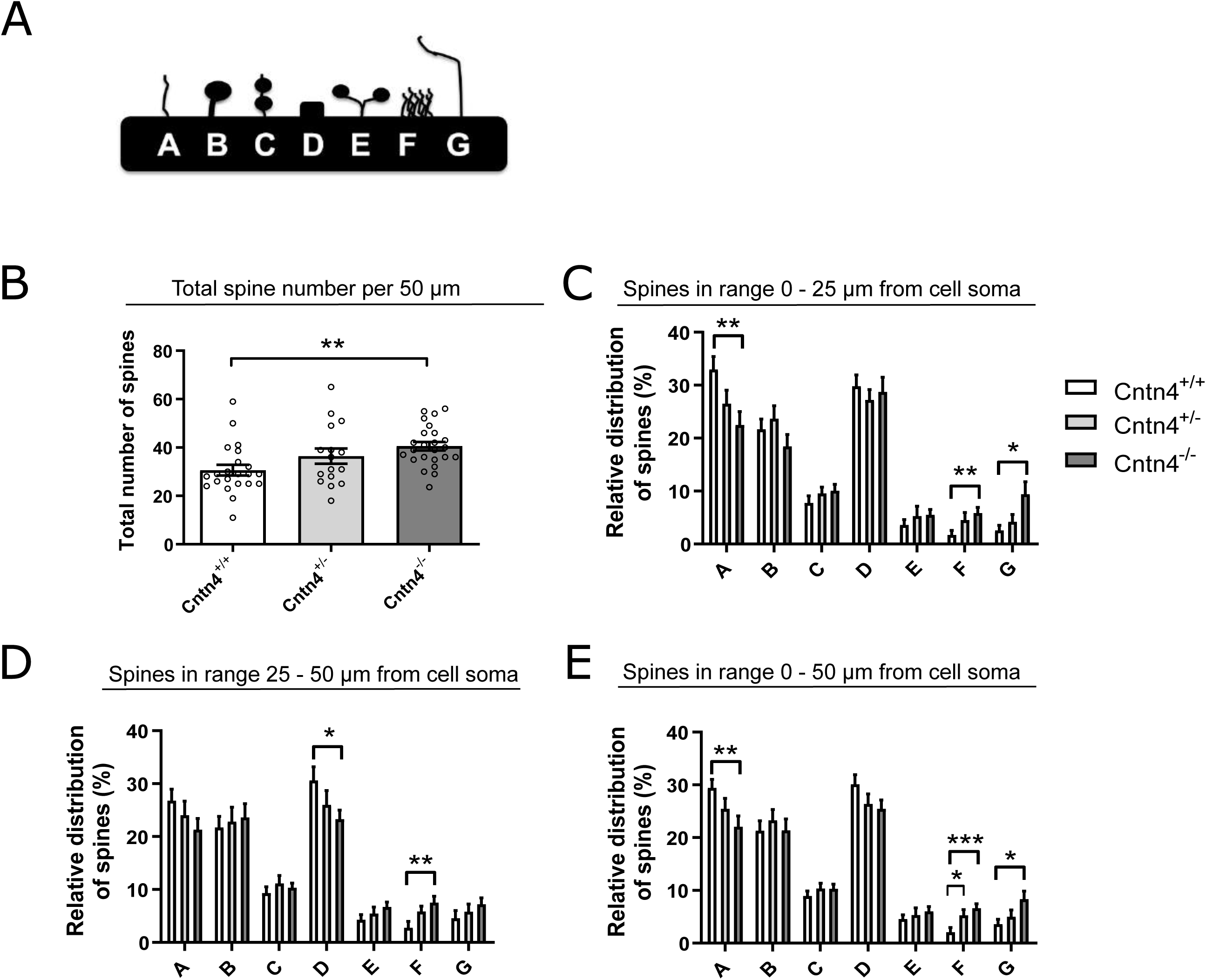
Spine analysis of M1 region, layers II-III pyramidal cells of adult *Cntn4^+/+^*, *Cntn4^+/-^* and *Cntn4^-/-^* mice. A) Schematic representation of morphology of spine maturation. Quantitative analysis reveals the B) total number of spines and relative expression of the maturation state of spines in C) first 25 µm; D) final 25 µm and E) total 50 µm. Analysis was performed on at least 5 mice for neurons, n=22 (*Cntn4^+/+^*), n=16 (*Cntn4^+/-^*), n=24 (*Cntn4^-/-^*) using one-way ANOVA. Data are presented as mean ± S. E. M.

### CNTN4 overexpression leads to longer neurites in primary culture

*In vitro* primary cell culturing was conducted to evaluate the consequences of CNTN4 overexpression (OE) on neurite outgrowth and cell body volume. Transfection of cortical neurons showed a significantly increased longest length of the dendrites (p = 0.0415, unpaired Student’s *t* test), although the total length was not significantly different between empty vector (EV) and CNTN4 OE neurons (p > 0.05, unpaired Student’s *t* test) (Figure 4A-B). No other morphological changes were observed in the neurons. No significant change in apoptosis was observed for CNTN4 overexpression (p > 0.05, unpaired Student’s *t* test) (Figure 4C).

**Figure 4:**
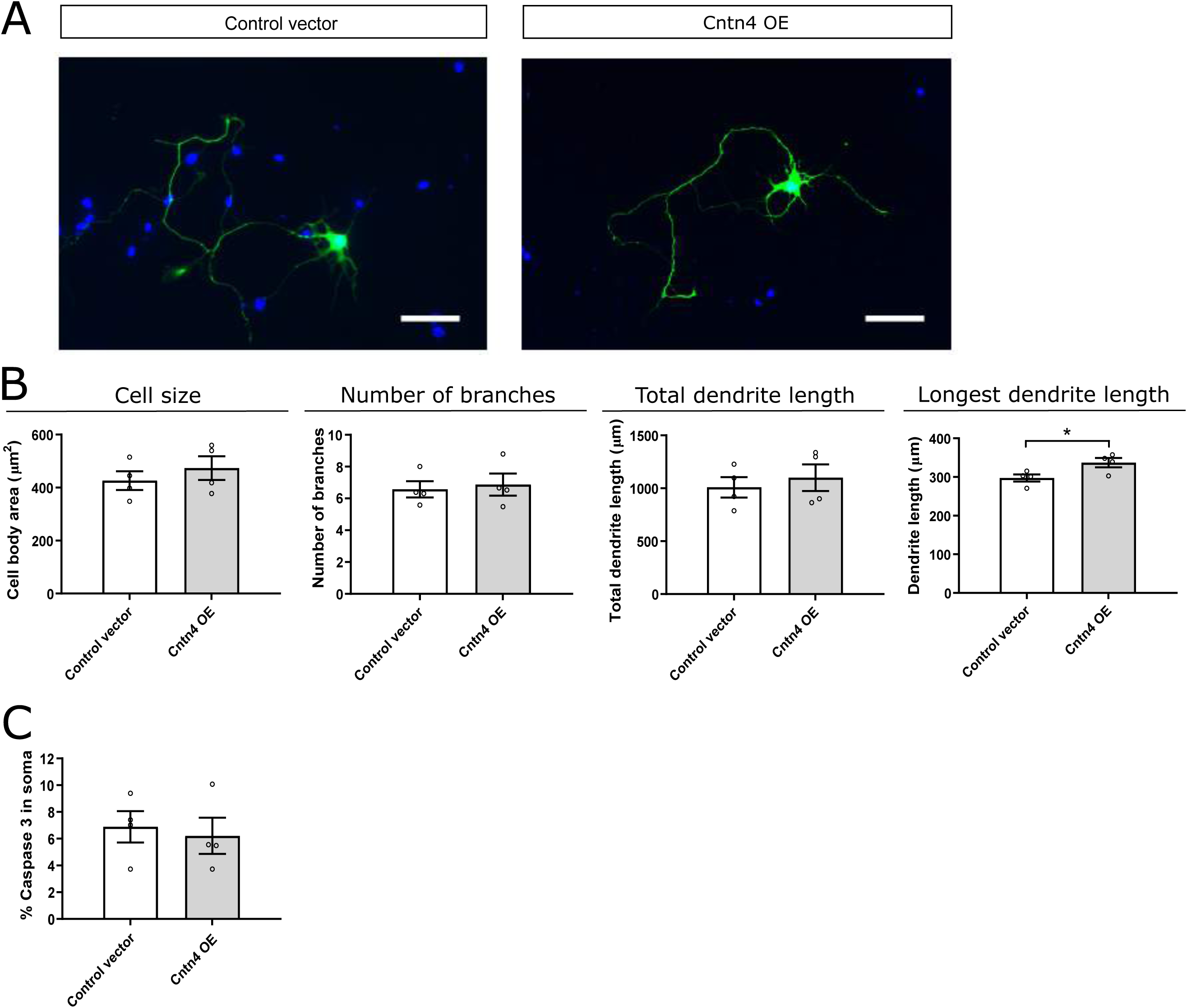
Primary culture of postnatal cortical neurons transfected with Cntn4 or an empty vector after 48 hours. Cortical cultures were prepared from mouse cortices at age P0-P1. Neurons were transfected with a control vector or CNTN4 on DIV2. A) Representative images of morphology of CNTN4 OE and empty vector transfected neurons. B) Average cell body surface in CNTN4 OE and empty vector transfected neurons; total number of dendritic branches; total length of dendritic branches and length of the longest dendrite (p = 0.0415, unpaired Student’s *t* test). Data are presented as mean ± S. E. M, four batches, total analysed cell number n=439 (Cntn4^+/+^), n=279 (empty vector). The scale bar represents 50 µm. C) Quantification of the percentage of Caspase-3 positive neurons shows no significant difference between the control and CNTN4 overexpression (p > 0.05, unpaired Student’s *t* test, n=427 total neurons analysed).

### Identification of APP as interacting partner of Cntn4

In order to find proteins that interact with CNTN4 to exert its role, an unbiased proteomics approach using CNTN4 protein fused to the Plexin A2 transmembrane domain and GFP (CNTN4-TMGFPBio and control TMGFPBio) was used to identify binding partners of this protein after expression in HEK293 cells (Figure 5A-B). These cells are known to express a large range of proteins (46), and have successfully been used before to find partners of other contactins (47). Following immunoprecipitation (IP) experiments, CNTN4 fusions and control proteins were detected at 148 and 40 kDa, respectively, by Western blotting and Coomassie blue staining (Figure 5C-D). Raw mass spectrometry data was analysed with the Mascot search engine and scores were assigned to identify peptides. In comparison to control experiments, confidence scores using Saint scoring (48) were assigned to the identified proteins. A ranked list of putative interacting proteins was obtained representing proteins that were significantly higher or exclusively present in the Cntn4 fusion protein pull-down samples (Table 1). The highest scoring transmembrane proteins were the proteins radixin (RDX), amyloid-beta precursor protein (APP) and brain acid soluble protein 1 (BASP1).

**Figure 5:**
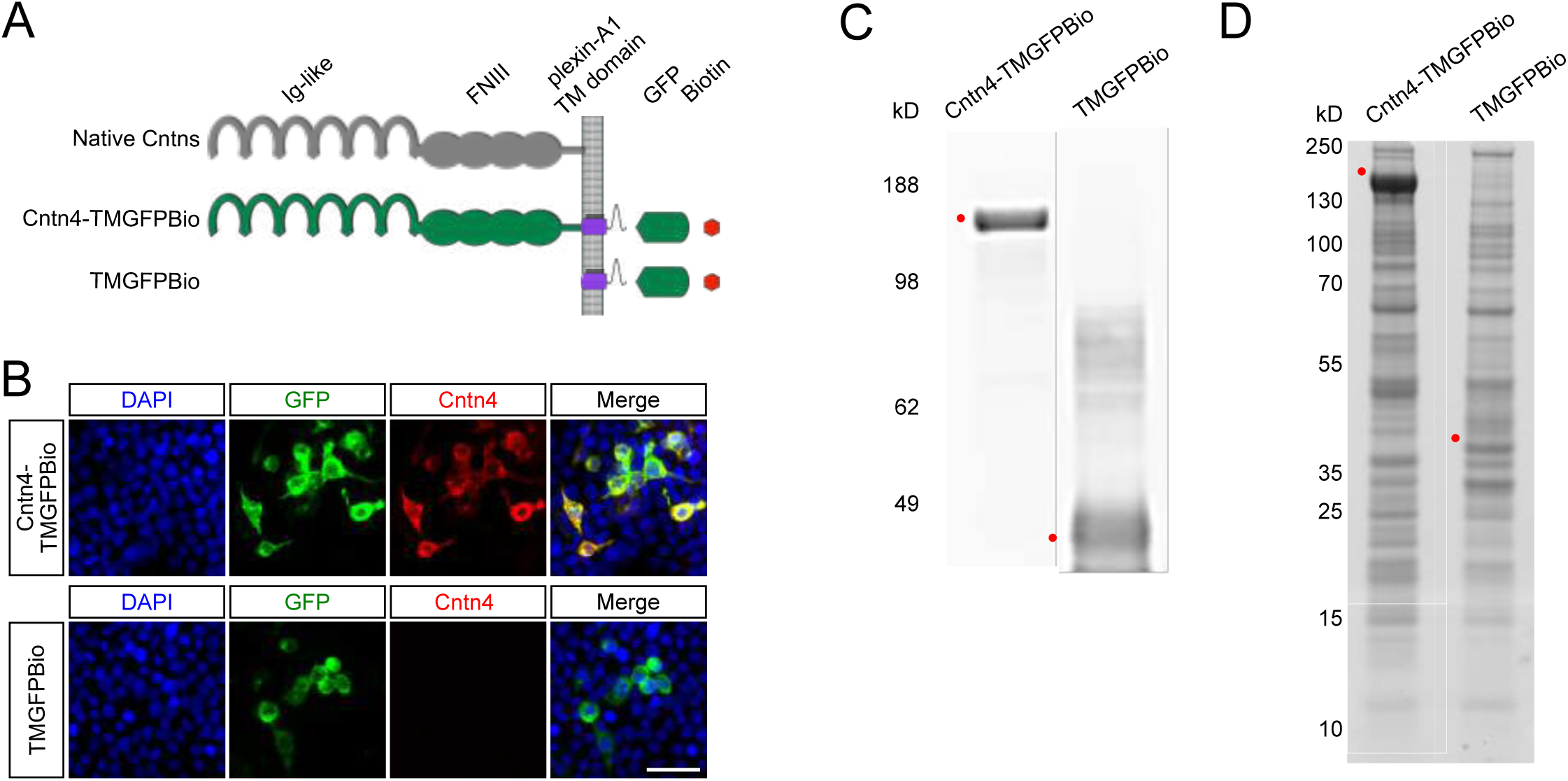
Identification of CNTN4 binding partners by proteomics. A) The architecture of the native CNTN4 protein and the structures of the CNTN4 and control fusion proteins tagged with GFP and biotin. B) Expression of CNTN4-TMGFPBio and TMGFPBio in HEK293 cells, detected by fluorescence (green), anti-CNTN4 antibody (red) and DAPI (blue) staining. Scale bar represents 30 µm. C) Precipitations were performed by anti-GFP-coupled beads and eluates from anti-GFP-coupled beads were analysed on Western blot using an anti-GFP antibody. D) Coomassie blue stained the NuPage 4-12% gels, which were submitted to mass spectrometry analysis. Red dots indicate respective expressed fusion proteins. Molecular weights are as follows: CNTN4-TMGFPBio = 147.8 kDa; TMGFPBio = 39.6 kDa. Ig-like, immunoglobulin-like; FNIII, fibronectin type III; TM, transmembrane domain; GFP, green fluorescent protein; Bio, biotin.

**Table 1:**
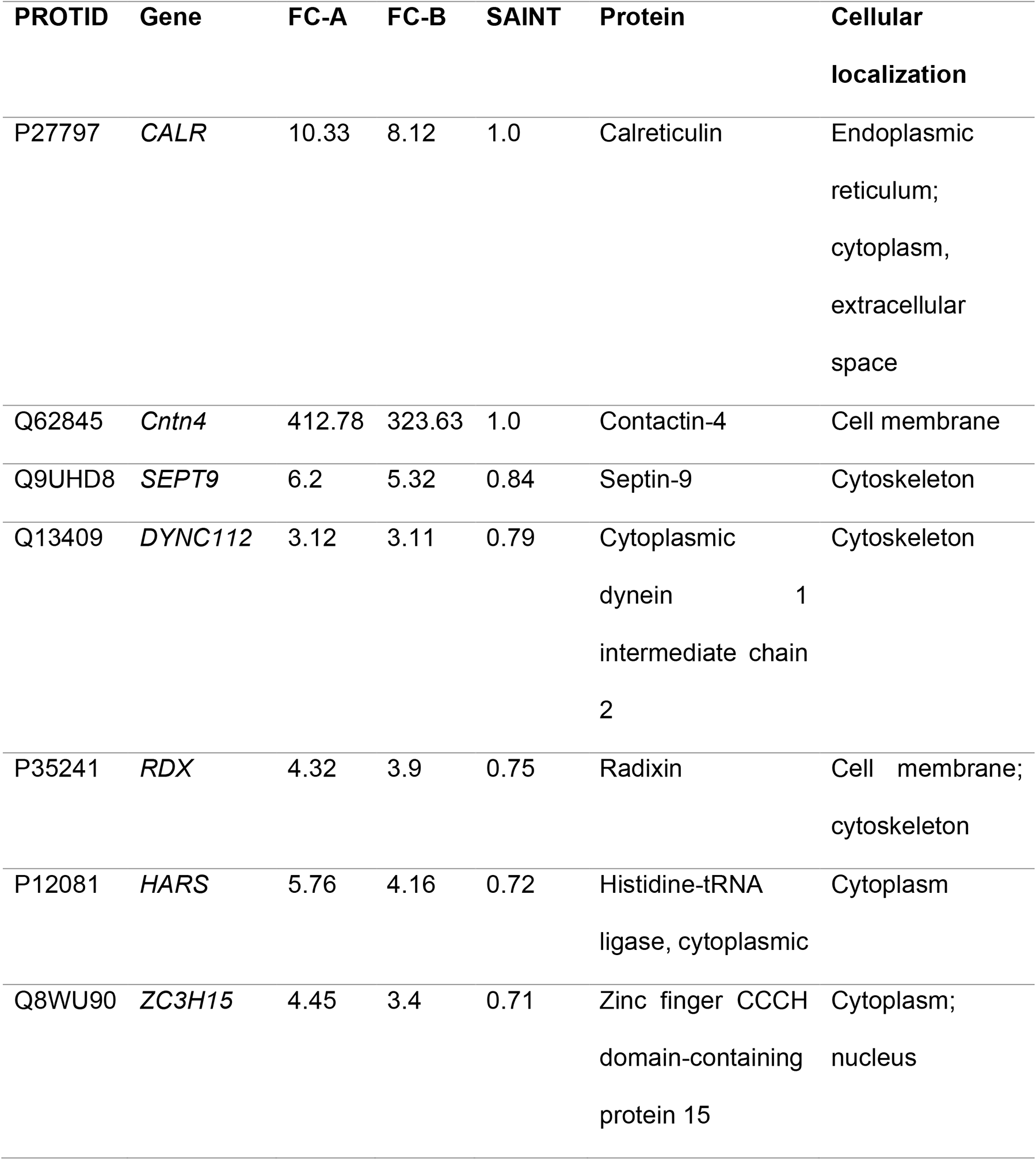
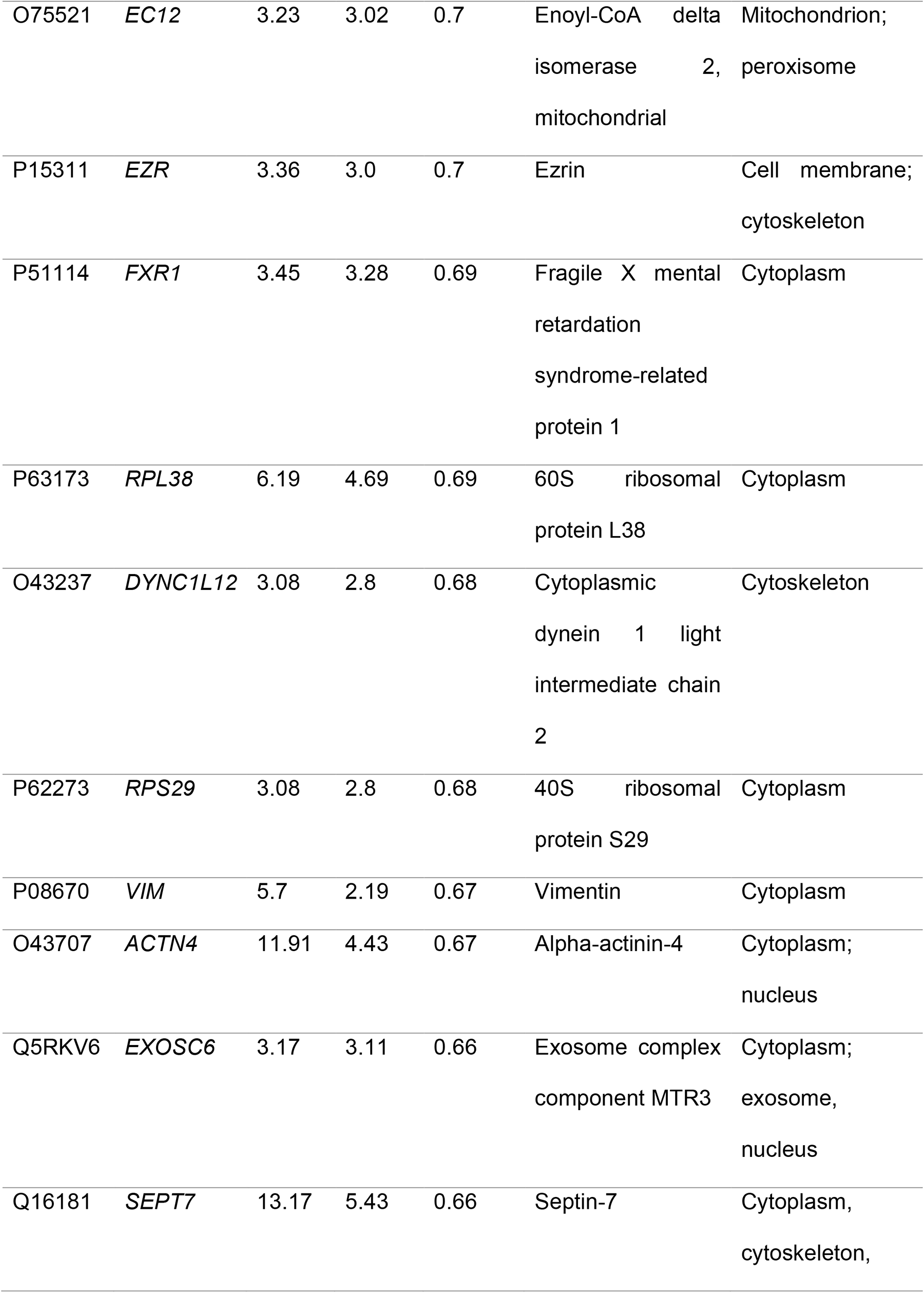
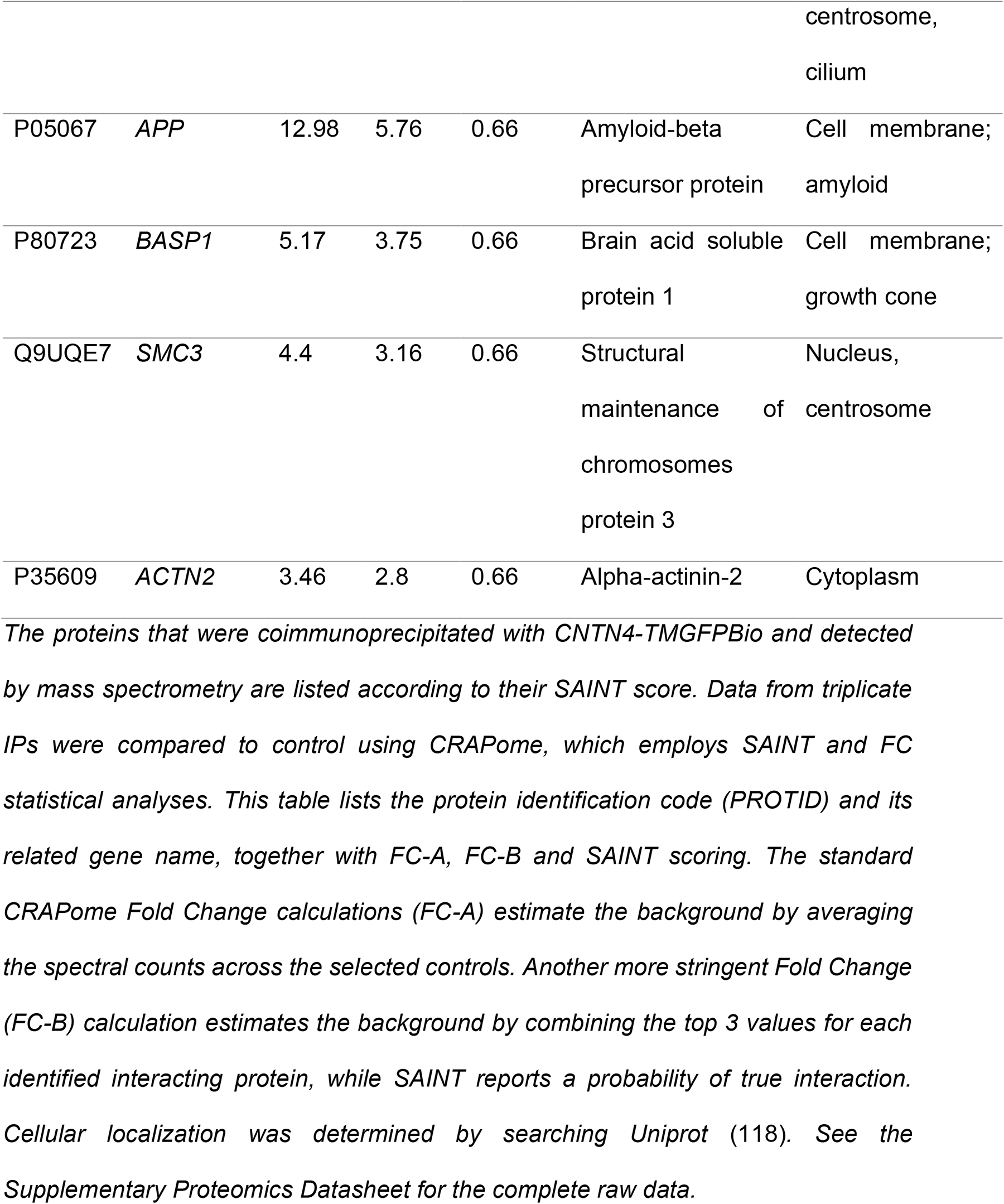
Top CNTN4 interacting proteins.

### Binding and *cis/trans* interactions between CNTN4 and APP

The SAINT and FC probability scoring confirmed APP as an interactor of CNTN4 (which has been previously described (49)), which we further examined *in vitro*. To validate the association of CNTN4 and APP, HEK293 cells were transfected with empty or native CNTN4 expression plasmids. IP analysis by Western blotting demonstrated coprecipitation of CNTN4 and APP, but not with control (Figure 6A). Investigating the cellular localisation of the two proteins in cell lines and neuronal cultures further supported the interaction of CNTN4 and APP. Native CNTN4 transfected in HEK293 cells and endogenous APP colocalised on the membrane of the same cell (Figure 6B), although the majority of APP was localised intracellularly. Similarly, immunostaining of cultured cortical neurons with anti-CNTN4 and anti-APP antibodies revealed endogenous localisation of these proteins throughout the neurons (Figure 6C). CNTN4 and APP were not clearly delineated on membranes of neurons and the bulk of the proteins seemed to localise intracellularly. Along the neurites both CNTN4 and APP were expressed in puncta, of which some overlap. These results showed that CNTN4 and APP partly colocalise on the cell membrane when co-expressed. To study whether CNTN4 and APP interact in *cis* or *trans*, cell adhesion assays were performed (Figure 6D). Separate populations of HEK293 cells were co-transfected either with CNTN4 together with DsRed or with APP together with EGFP expression plasmids. As a positive control, cells were co-transfected with NLGN1 and DsRed or with NRXN1β^-^ and EGFP expression plasmids. NLGN1 and NRXN1β^-^ are well-established *trans*-binding partners (50,51). Since CNTN4 can form homodimers in *trans* (52), cells were also co-transfected with CNTN4 and EGFP or Cntn4 with DsRed expression plasmids, as a second positive control. As a negative control, cells were transfected with either DsRed or EGFP expression plasmids. Cells expressing DsRed, NRXN1β^-^ or APP, were incubated with cells expressing EGFP, NLGN1 or CNTN4. As a negative control, cells expressing DsRed were incubated with cells expressing EGFP. Cell aggregation was measured and quantified after incubating the cell mixtures for up to 90 min (Figure 6E). A significant number of adhering cell clumps was observed when NRXN1β^-^-expressing cells (green) were incubated with NLGN1-expressing cells (red), serving as a positive control. A significant degree of cell-aggregation was also observed in the mixture of CNTN4-expressing cells (green) with CNTN4-expressing cells (red), demonstrating CNTN4’s capability homodimerization (52). It was observed that CNTN4-expressing cells (red) aggregated with APP-expressing cells (green) to a similar degree as the positive interaction between NRXN1β^-^-and NLGN1-expressing cells. All three test conditions showed significant aggregation compared to the negative control. These data demonstrate that the binding of CNTN4 and APP can occur when the proteins are expressed on opposing cells in *trans* configuration but cannot rule out *cis* interaction as well. Subsequently the cell surface binding assay (47) was performed which confirmed the *cis* biochemical interaction between CNTN4 and APP. The binding between soluble, tagged APP-GFP and membrane-bound FLAG-CNTN4 was measured in HEK293 cells via confocal microscopy (lower panel, Figure 6G). The well-characterised *trans*-interacting proteins neogenin and RGMa were used as positive controls in this assay (top panel, Figure 6G). Quantification shows a significant number of double labelled transfected cells in the condition of CNTN4 and APP (Figure 6H), which confirms a *cis* interaction between CNTN4 and APP.

**Figure 6:**
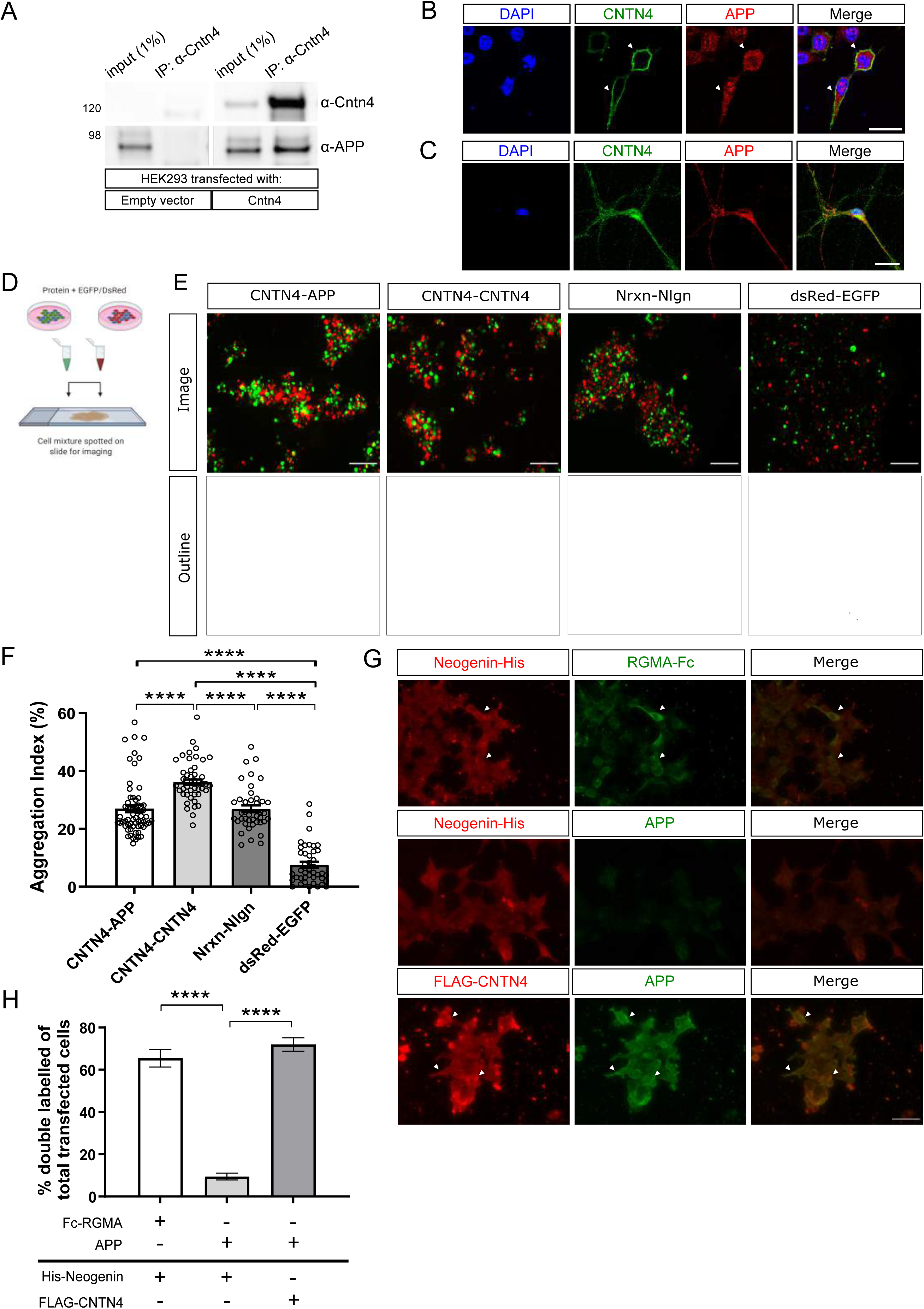
Interaction of CNTN4 and APP. HEK293 cells were transfected with empty vector or CNTN4 expression plasmid, coding for the native Cntn4 protein. A) After IP of the CNTN4 by an anti-CNTN4 antibody the eluates were analysed by Western blot. Blots stained with antibodies against CNTN4 and APP revealed interaction between CNTN4 and APP. No coprecipitation was found in normal IgG control IP. B) HEK293 cells expressing CNTN4 protein (green) colocalised with endogenous APP (red) on the membrane. Arrows indicate co-expression on the membrane. DAPI staining is in blue. The scale bar represents 20 µm. C) Endogenous CNTN4 (green) and APP (red) are co-expressed in neurons. DAPI staining is in blue. The scale bar represents 20 µm. D) Schematic overview of the cell adhesion assay, in which populations of HEK293 cells were co-transfected with different proteins and either EGFP or DsRed expression plasmids. Two HEK293 cells populations were combined and incubated. Cell suspensions were spotted onto slides for imaging by fluorescence microscopy. Created with BioRender.com. E) Cells expressing either EGFP alone or together with NRXN1β^-^ or APP (green) were mixed with cells expressing DsRed alone or together with NLGN1 or CNTN4 (red). Aggregation of cells expressing NRXN1β^-^ + EGFP with NLGN1 + DsRed, CNTN4 + EGFP with CNTN4 + DsRed, and CNTN4 + DsRed with APP + EGFP was observed. There was no aggregation of cells expressing EGFP + DsRed. The scale bar represents 200 µm. F) Aggregation index was determined from five fields of 1.509 mm^2^ per cell suspension combination of each independent cell adhesion assay (n=3). Analysis was performed using one-way ANOVA. The graph bars are presented as mean ± SEM. ****p< 0.0001. G) HEK293 cells expressing Neogenin-His or FLAG-CNTN4 were incubated with soluble ecto-domains of RGMA-Fc and APP. Upper panel: Neogenin-His-expressing cells (red) bound soluble RGMA-Fc (green), but not with APP (green) (middle panel). Lower panel: FLAG-CNTN4-expressing cells (red) bound soluble APP (green). Scale bar represents 50 µm. Arrowheads indicate examples of green and red overlay. H) Quantification of about 300 transfected cells per transfection condition of each independent cell surface binding assay (n=3) was performed. Statistical analysis was performed using unpaired Student’s *t* test and one-way ANOVA. The graph bars are presented as mean ± SEM. ****p<0.0001.

### APP deficiency causes abnormal apical dendrite morphology

Since the CNTN4-APP interaction suggested an involvement of APP in the pyramidal neuron phenotypes of cortical neurons, we investigated the morphology of Golgi-stained pyramidal neurons in *App^+/+^*, and *App^-/-^* mice (Figure 7A-B). Pyramidal neurons in the motor cortex layers II-III were analysed for dendrite length, branching and complexity. There were no significant differences in apical or basal neurite length, DCI, total intersection number and last Sholl intersection (Figure 7C, E). However, there were significant morphological changes of the apical dendrite, since the number of apical dendrite tips was significantly decreased in the *App^-/-^* mice compared to the *App^+/+^* mice (p = 0.0276, *App^+/+^* vs *App^-/-^*, unpaired Student’s *t* test) (Figure 7C).

**Figure 7:**
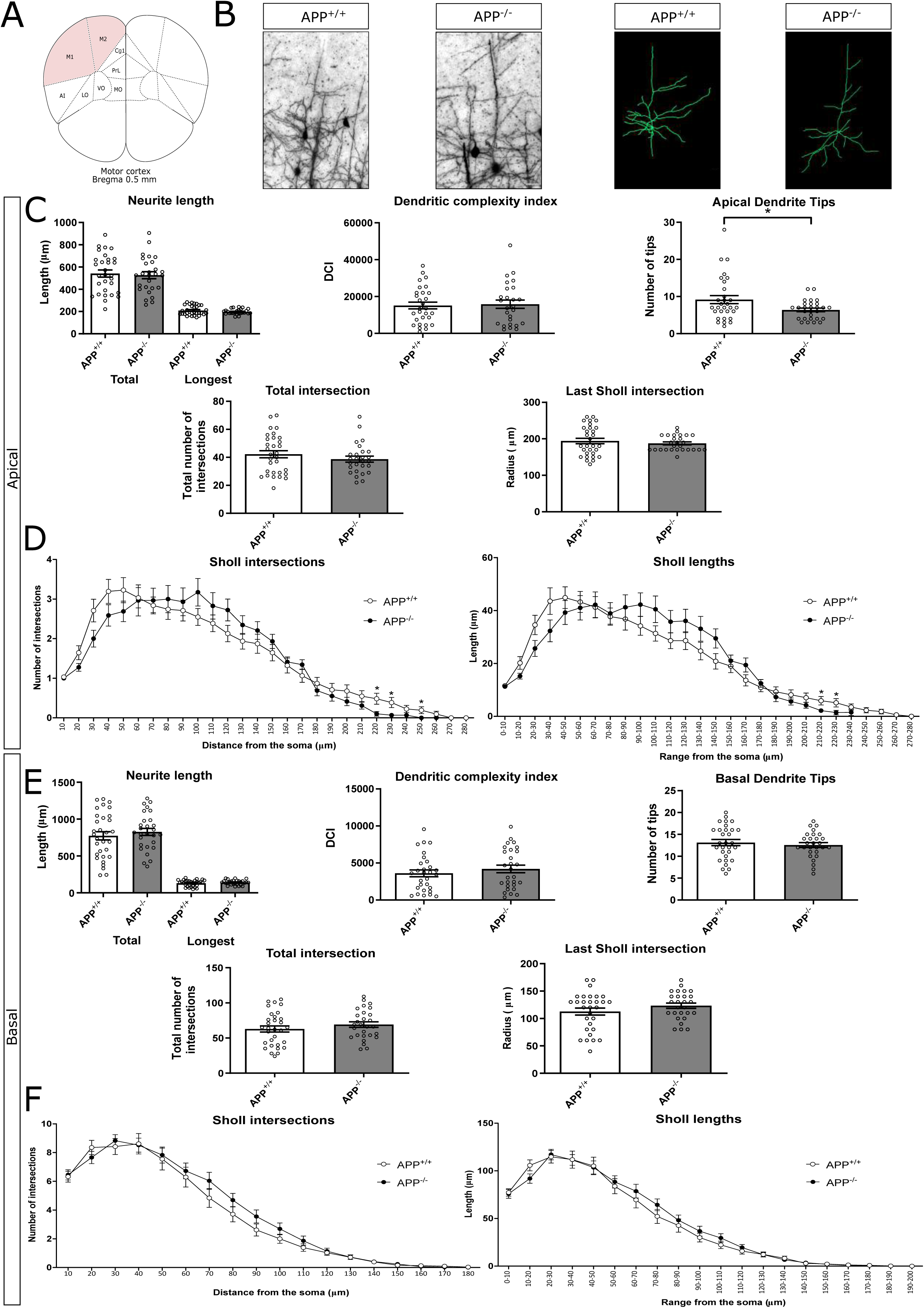
Neuron morphology analysis results for the primary motor cortex. A) Schematic representation of the motor cortex with labelled Bregma anterior-posterior. Adapted from Paxinos and Franklin, 2001. B) Golgi staining in *App^+/+^* and *App^-/-^* mice motor cortex and trace outlines of example pyramidal neurons. The scale bar represents 40 µm. C) & E) Quantitative morphological results for the apical and basal dendrites respectively. D) & F) Sholl plots indicate the distribution of respective apical and basal dendritic intersections and length at increasing distance from the centre of the cell body. Quantitative analysis was performed, in each area, on at least six slices in *App^+/+^* and *App^-/-^* mice (n=5 per genotype). Data are presented as mean ± S. E. M, cell number n=30 (*App^+/+^*), n=30 (APP^-/-^). (p = 0.0276, *App^+/+^* vs *App^-/-^*, apical dendrite tips). Statistical analyses were carried out using unpaired Student’s *t* test.

Sholl plots indicated the distribution of dendritic intersections and length at increasing distance from the centre of the cell body. The *App^-/-^* mice had significantly fewer Sholl apical intersections in the range of 220-250 µm from the soma, compared to *App^+/+^* mice. Furthermore, significantly shorter Sholl apical lengths in the range of 210-230 µm from the soma were observed in *App^-/-^* mice (Figure 7D). Taken together, the morphological changes in pyramidal neurons, as observed in *Cntn4^-/-^*, were not observed in *App^-/-^*.

### Expression levels and morphology of CNTN4-and APP-deficient human cells

The neuroblastoma cell line SH-SY5Y is a well-established model for differentiation of cells into cortical-like neurons (53,54), and was used to evaluate the single and double loss-of-function of CNTN4 and APP on neuronal differentiation *in vitro*. *CNTN4^-/-^, APP^-/-^,* and *CNTN4^-/-^/APP^-/-^* SH-SY5Y neuroblastoma cell lines were generated by CRISPR-Cas9 gene editing (Figure 8A-C and Supplementary Materials). A cell line created by transfection of empty CRISPR vector was used as a control cell line (EV). qPCR was performed on all knockout cells with primers designed for targeting *CNTN4* and *APP* (Figure 8D-E). Expression levels of *Cntn4* were found to be significantly reduced by approximately 50% in the *APP^-/-^* SH-SY5Y cell line, relative to EV expression levels. Conversely, expression levels of *APP* were reduced by approximately 50% in the *CNTN4^-/-^* and EV SH-SY5Y cells. As expected, both *CNTN4* and *APP* expression levels were reduced in the *CNTN4^-/-^/APP^-/-^* SH-SY5Y cells.

**Figure 8:**
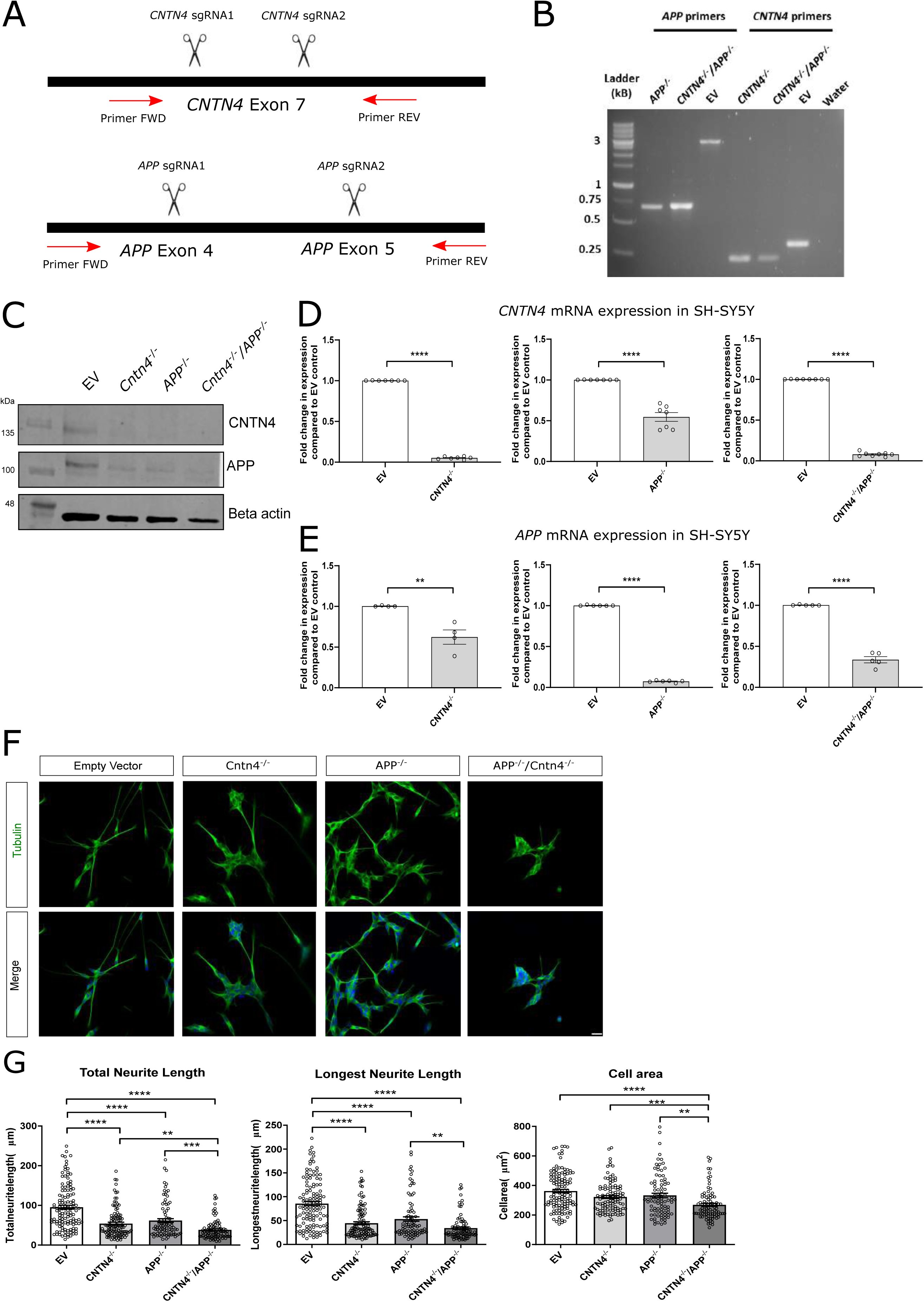
CNTN4 knockout cell lines show morphological differences and significantly higher levels of apoptotic activity. A) Schematic representation of the sgRNA designed to cause an indel in *CNTN4* and *APP*, respectively. B) SH-SY5Y clonal cells were screened via PCR using specific primers for *CNTN4* and *APP.* SKO = single knockout, DKO = double knockout, EV = empty vector control, W = water control. Expected sizes were for APP EV = 2.905 kB, APP knockout = 0.701 kB, CNTN4 EV = 0.341 kB, CNTN4 knockout = 0.249 kB. 1kB ladder (Solis Biodyne). C) Protein extracted from SH-SY5Y cells was analyzed by Western blot. Blots stained with anti-CNTN4 and anti-APP antibodies revealed lower expression in *CNTN4^-/-^, APP^-/-^, CNTN4^-/-^/APP^-/-^* cells, respectively. Western blotting was performed on four SH-SY5Y cell lines (empty vector, *CNTN4^-/-^, APP^-/-^, CNTN4^-/-^/APP^-/-^*). Molecular weights are as follows: CNTN4 = 150 kDa; APP = 100-140 kDa; beta-actin = 47 kDa. D) Fold change of *CNTN4* and E) *APP* expression levels in *CNTN4^-/-^, APP^-/-^, CNTN4^-/-^/APP^-/-^* cells compared to empty vector cells, generated by qRT-PCR. Analysis was performed on three SH-SY5Y cell lines (*CNTN4^-/-^, APP^-/-^, CNTN4^-/-^/APP^-/-^*) using one-way ANOVA and Tukey’s multiple comparison post-hoc test. Primer sets *Cntn4* P2 and *App* P4 are presented, but two further primer sets are presented in Figure S2. All primer detail is included in the Supplementary Materials. F) Representative images of morphology of empty vector, *CNTN4^-/-^*, *APP^-/-^* and *CNTN4^-/-^/APP^-/-^* cells. Tubulin staining is in green and DAPI staining is in blue. The scale bar represents 25 µm. G) Quantitative morphological analysis was performed on at least 3 coverslips in empty vector, *CNTN4^-/-^*, *APP^-/-^* and *CNTN4^-/-^/APP^-/-^* cells (n=3 per genotype). Data are presented as mean ± S. E. M, cell number n=30 per genotype. * p < 0.05. Statistical analyses were carried out using unpaired Student’s *t* test.

Knockout SH-SY5Y cells were differentiated using retinoic acid, BDNF and lack of FBS over a five day period, adapted from Shipley *et al.* (55). Morphological analysis was carried out on differentiated cells. It was observed that the total and longest neurite length was significantly reduced in the *CNTN4^-/-^ APP^-/-^* and *CNTN4^-/-^/APP^-/-^* cells compared to the EV cells (Figure 8F-G). It was further observed that the cell area was significantly reduced in the *CNTN4^-/-^, APP^-/-^* and *CNTN4^-/-^/APP^-/-^* cells compared to the EV cells (Figure 8F-G). This suggests that the interaction between CNTN4 and APP contributes to their gene expression and contribution to neurite outgrowth.

## Discussion

We have shown that *Cntn4*-deficient mice have abnormalities in the motor cortex. This is important since CNTN4 has previously been associated with neurodevelopmental disorders such as ASD (28,36,37). To understand the aetiology of disorders such as ASD, and the role that CNTN4 has, we set out to examine the pathological effects of loss-of-function of this Contactin family member. The loss of *Cntn4* does not affect gross anatomical development, such as body weight and brain size in mice (44).

Here we showed that that *Cntn4*-deficient mice have significantly reduced cortical thickness in the M1 motor cortex region and altered dendrite length and complexity in M1 neurons. We interpret that CNTN4 mildly affects the numbers of pyramidal neurons in M1. No other cortical areas other than the motor region were shown to be affected in *Cntn4-*deficient mice (Figure S1), suggesting that CNTN4 has a specific role for the M1 region. The motor cortex is in close proximity to the main olfactory bulb and accessory olfactory bulbs, and these respective regions have shown CNTN4 to be an axon guidance molecule that mediates neuronal wiring (56). Kaneko-Goto *et al.* demonstrated the importance of CNTN4 in organising the wiring of the olfactory system, where CNTN4 deficiency leads to aberrant connections of the neurons to the olfactory receptors (41). Our research revealed comparable findings in the M1 region since *Cntn4-*deficient mice revealed abnormal cell outgrowth and arborisation in proximity to the cell soma, and increased dendrite length (Figure 2). This suggests a role for CNTN4 in dendrite outgrowth and directionality in M1 motor cortex.

Disrupted neuron migration, abnormal neurite growth and spine formation in the cortex are all associated with neurodevelopmental disorders such as ASD (57,58). Nissl and Golgi staining, combined with cortical neuron morphological studies, all point towards a role for CNTN4 in neurite outgrowth. *Cntn4*-deficient mice demonstrate decreased cortical thickness in the motor region, abnormal spine density and decreased length of the longest neurite (Figures 1-3). The latter is also in agreement with human *CNTN4^-/-^* SH-SY5Y cell morphology (Figure 8G). Moreover, overexpression of CNTN4 in cortical neurons causes an increased length of the longest neurite but no significant change in apoptosis (Figure 4). These morphology and apoptosis results agree with previous reports of CNTN6 overexpression in cortical neurons (47) and CNTN4-6 overexpression co-culture experiments of rat cortical neurons and HEK293 cells (59). Cell adhesion molecules such as CNTNAP2 and Contactin family members are known to play crucial roles in neuritogenesis (59,60), and deficits such as abnormal cortical migration in the *Cntnap2^-/-^* mouse model have been linked to ASD (61).

Spines contain a range of proteins amongst which receptors, cytoskeletal and adaptor proteins, and associated signalling molecules (62) are dynamic in response to synaptic activity (63). Kaneko-Goto *et al.* identified that CNTN4 expression is regulated by neuronal activity (56) and Zhao *et al.* found enriched expression of CNTN4 at excitatory synapses on dendritic spines (64). *Cntn4*-deficient mice showed an increased spine density in cortical neurons (Figure 3), suggesting CNTN4 is not required for the formation of cortical spines. Interestingly, Zhao *et al.* observed that knockdown of CNTN4 by shRNA transfection reduced the spine density of primary cortical neurons in culture (64), whereas the present results were the opposite of these findings. This suggests some brain area-specificity in the mouse brain may be altering the cascade that contributes to the formation of spines. In further contrast, the spine density in hippocampal neurons decreased with *Cntn4* deficiency (44), pointing towards a cell type specific role for CNTN4. It was further observed that there were less immature and more abnormal cortical spines found in the absence of *Cntn4,* suggesting an important role in synapse quality. This suggests that while *Cntn4* contributes to spine formation in the cortex and hippocampus, it has an alternate role in the spine formation process. Spine maturity is suggested to be the subject of glutamate receptor input (63,65) and, as a consequence, influences the excitation-inhibition balance. Previous reports observed that a disruption in CNTN4 causes a decrease in number of excitatory synapses and therefore a decrease in neural activity (64). Similar findings have been further identified for other cell adhesion proteins such as neuroligin (NLGN) and protocadherin 9 (PCDH9) (66–70). Synaptic expression and localisation of CNTN4 overlaps with synaptogenesis in the developing brain (69,71). This previous research links CNTN4 to motor development and suggests a broader role in the excitation-inhibition balance via spine maturation. Our data also suggests a role for *Cntn4* in the maturation of spine development, which may influence the excitation-inhibition balance, and contribute to the stability of the synapse after formation. The role of dysregulated CNTN and CNTNAP at the synapse has been previously postulated (72).

The question is whether CNTN4 deficiency alone is causal of the abnormalities or that CNTN4 is part of a crucial interaction. To answer this, binding assays were carried out to reveal the interacting partners of CNTN4 and understand their function. Osterfield *et al.* (49) showed that CNTN4 and APP were direct binding partners, which we confirm through our independent unbiased proteomics screen (Figure 5 & Table 1) and binding assays (Figure 6). In this study CNTN4 appeared to be bound to full length APP since the tryptic peptides found in mass spectrometry were derived from full length APP and co-immunoprecipitation validated this interaction between CNTN4 and APP (Figure 6). APP is known to involved in *cis* and *transcellular* adhesion (73) and is a synaptic adhesion molecule (74,75). CNTN4 and APP may stabilise one another at the cell surface, which is subsequently important for synaptogenesis.

Osterhout *et al.* observed that in the absence of CNTN4 or its binding partner APP, deficiencies were observed in the developing chick retinotectal system (42,76). APP, a transmembrane protein, is unquestionably associated with Alzheimer’s disease but also serves crucial physiological functions (77) for neuronal migration, neuronal morphology, synaptic plasticity, learning and memory (78) and more recently also for ASD-like abnormalities (79). The interaction between Contactin family members, including CNTN4, and APP has been reviewed elsewhere (72). In particular, APP is highly expressed in the developing cortex during differentiation and migration of cortical neurons (80–82). Osterfield *et al.* demonstrated that the soluble isoform of CNTN4 is secreted by neurons, and creates a gradient upon which dendrites can grow (49). This agrees with our finding that overexpression of CNTN4 in primary neurons resulted in significantly longer dendrites (Figure 4). Similarly, Osterfield *et al.* showed that APP also promoted neurite outgrowth (49). However, interestingly, the *App^-/-^* mice had a mild phenotype, and there was little evidence of morphological abnormality (Figure 7). More pronounced effects are observed in aged *App^-/-^* mice, in the spine density and dendritic branching of CA1 and LII-III neurons (83). Compensation by other family members such as APLP1 and APLP2 is discussed later, however, it is worth noting here that the phenotype of the combined mutants of the APP family is much more severe than the single APP KO (78,79). Furthermore, APP itself regulates many genes (84), and there are many genes which regulate the process of APP cleavage (85), and therefore, it would be of future benefit to investigate the genes responsible for regulating APP gene expression, for example the downregulation of two genes PSMA5 and PSMB7 involved in APP-induced cell proliferation impairment (86).

CNTN4 also interacts through its Ig-like domains with PTPRG, which is expressed primarily in the nervous system and mediates cell adhesion and signalling events during development (87,88). We did not further investigate this interaction since PTPRG was not identified in our proteomics screen. Additionally, Mercati *et al.* found differences in neurite outgrowth were not caused by the CNTN4 and PTPRG interaction (59). Candidate proteins identified to be APP binding partners include F-spondin (89), collagen, netrin-1, laminin, and the Aβ peptide (90). APP has been shown to have a direct high affinity interaction with contactin 3 and 4 (49). In functional assays of cultured retinal ganglion cells (RGCs), CNTN4 and APP modulated axon behaviour specifically in the context of NgCAM-dependent axon growth, demonstrating functional interactions among these proteins.

In order to investigate mediating effects, *in vitro* studies of CNTN4 and APP deficiency were carried out (Figure 8). CNTN4 mRNA and protein expression levels were significantly reduced in the *APP^-/-^* SH-SY5Y cell line, and vice versa APP mRNA and protein levels were significantly reduced in the *CNTN4^-/-^* SH-SY5Y cell line (Figure 8). It was evident from mRNA expression levels that CNTN4 and APP were dependent on one another, although we cannot say exclusively. Osterfield *et al.* demonstrated that CNTN4 had the highest binding affinity to APP (49), and we further observe that the interaction has approximately 50% impact on mRNA and protein levels. This suggests that a complex needs to be formed for mutual stability, and furthermore, the binding of CNTN4-APP can be functionally meaningful at around 50% of their protein levels. In addition, there were significant morphological deficiencies in all knockout SH-SY5Y cell lines compared to the empty vector control. The most severe morphological deficiencies were observed in the *CNTN4^-/-^/APP^-/-^* SH-SY5Y cell line, which further emphasises the importance of the CNTN4 and APP interplay.

The FnIII domain of CNTN4 is reported to interact with the E1 domain of APP, and other APP family members, on the cell surface (49,91). Zhao *et al.* observed CNTN4 proteins lacking FnIII or GPI domains lack the ability to regulate dendritic spine formation (64). Our hypothesis is that when CNTN4 is deficient, 1) the function to which the binding of CNTN4 to APP contributes is lost, and the ability of CNTN4 to regulate dendritic spine formation diminishes which would cause abnormal neurite outgrowth; 2) the loss of CNTN4 would cause other proteins to alternately bind to the E1 domain of APP and affect neurite outgrowth. These two possibilities were considered. The results of the present study demonstrate the first possibility (Figure 8), but further studies on the second are needed. There is also the important consideration of the amyloidogenic and non-amyloidogenic APP processing pathways, and whether the lack of CNTN4 interaction has an effect. For example, APPsα is a cleaved ectodomain of APP, which has been shown to modulate cell behaviours including neurite outgrowth, synaptogenesis, neurogenesis, and cell survival and proliferation (92–94). In this capacity, the interaction of CNTN4 with APP, may have potential therapeutic effects. In the case of APP deficiency, CNTN4 is hypothesised to (more highly) bind to other APP family members, such as APLP1 or APLP2 which conserve the E1 domain (95,96). This in turn offers further therapeutic benefits since APLP1 and APLP2 cannot generate an amyloidogenic fragment since they lack an internal Aβ site. However, the physiological function of the other APP family members is not fully understood. Missense mutations in the APP gene have been shown to cause familial AD (97). The physiological relationship between APP and Contactin members and their role in pathophysiology of AD is reviewed elsewhere (72). Our data suggests the interplay between CNTN4 and APP is important for synapse maintenance and neuronal stabilisation.

This research identified structural abnormalities in the brains of *Cntn4^-/-^* mice. Reduced cortical thickness of the layers of the M1 region was identified, which echoes connectivity and phenotypic abnormalities observed in ASD patients. Brimberg *et al.* observed cortical thinning in male mice after administration of Contactin-associated protein Caspr2-reactive antibody cloned from the mother of an ASD child (98), which aligns with the gene-dosage dependent cortical layer thinning in the motor cortex. Sensorimotor development impairments and ASD stereotypic motor movements have been previously observed in Pcdh9 and Cntnap2 knockout mouse models, respectively (61,99). Motor abnormalities correlate with ASD severity and are persistent in child development as well as in adults clinically diagnosed with ASD (100–104). Moreover, functional studies in human ASD patients suggest that subnetworks in the M1 region are alternately activated in comparison to normal development and within equivalent age groups showed that motor impairments are caused by underlying structural abnormalities or abnormal connectivity within brain networks in M1 cortex (105–108).

To summarise, our study identified differences in cell density and differentiation in the M1 region of CNTN4 deficient mice and a role for CNTN4 in neurite outgrowth and directionality. The interaction between CNTN4 and APP was also shown to be important in knockout human cell lines, with significant defects in cell morphology and elongation and the observation that CNTN4 mediates neurite outgrowth by binding to APP. This provides an important step forward in our understanding of Alzheimer’s disease, ASD and this field of neuroscience. It will be important for future neuropsychiatric studies to clarify how CNTN4 affects APP binding and how it is involved in the APP process.

In conclusion, our study revealed variations in cell density and differentiation in the M1 region of CNTN4-deficient mice, along with CNTN4’s role in guiding neurite growth and direction. The significance of CNTN4’s interaction with APP was also confirmed in human cell lines lacking CNTN4, highlighting defects in cell structure and elongation. Notably, our findings suggest that CNTN4 facilitates neurite extension through its interaction with APP. This advancement enhances our comprehension of Alzheimer’s disease, ASD, and the broader neuroscience field. For forthcoming neuropsychiatric investigations, elucidating CNTN4’s impact on APP binding and its involvement in APP processing will be pivotal.

## Materials and Methods

### Animals

*Cntn4*-deficient mice were kindly provided by Dr. Yoshihiro Yoshihara (RIKEN, Japan) (56). These mice were generated using a standard gene-targeting method as previously described. All experimental procedures were performed according to the institutional guidelines of the University Medical Center (UMC) Utrecht. All animal procedures were performed according to NIH guidelines and approved by the European Council Directive (86/609/EEC). For further generation and genotyping detail, see Supplementary Materials.

*App* knockout mice (*App^-/-^*) and littermate wild types (*App^+/+^*) were described previously (109). Fresh brain tissue extracted from male, five-month-old mice were prepared as described in Golgi staining section.

### Nissl staining

Brains were sectioned with a cryostat (Leica Microsystems, Wetzlar, Germany) coronally at 40 µm from rostral to caudal. Sections were mounted onto Superfrost slides (VWR, USA, 631-0108). The slices were rehydrated in graded levels of decreased concentrations of alcohol and then stained in 0.5% Cresyl-Violet (Sigma Aldrich, USA, 190-M) for 5 minutes. Finally, the slices were dehydrated in graded levels of increased concentrations of alcohol, cleaned in xylene, and then cover slipped using Entellan® (Merck, Damstadt, Germany, 107961).

Slices were imaged using light microscopy (Zeiss Axio Scope.A2, Germany) using the following stereotaxic coordinates (110): Bregma anterior-posterior +0.5 mm (frontal motor cortex), -1.70 mm (primary somatosensory cortex), and -2.80 mm (visual cortex).

Analysis of cortical thickness was performed within the frontal motor cortex (+0.5 mm to bregma), primary somatosensory cortex (-1.70 mm to bregma) and visual cortex (-2.80 mm to bregma) in all used brains. For cortical layer thickness, the superficial (layer I-IV) and deeper layers (layer V and VI) were measured, in each area, on at least three slices in *Cntn4*^+/+^ (n=7), *Cntn4*^+/-^ (n=4) and *Cntn4*^-/-^ mice (n=4) using ImageJ software (111).

### Immunohistochemistry

Brains were sectioned as described above and free-floating slices were stored in 0.02% sodium azide until immunohistochemistry was performed. The sections were washed with PBS and incubated in blocking buffer (1% BSA, 0.2% fish skin gelatin (Sigma Aldrich, G7765), 0.1% Triton X-100 in PBS) for 45 min. Sections were washed and incubated in permeabilization buffer (0.3% Triton X-100 in PBS) for 10 min before incubation with primary antibody in blocking buffer at 4°C for 2 hr. The sections were washed in PBS and pre-incubated with blocking buffer before incubating with secondary antibody at RT for 2 hr. The sections were embedded with Polyvinyl alcohol mounting medium with DABCO® anti-fading (Sigma Aldrich, 10981) onto glass slides after additional PBS wash steps.

Immunohistochemistry primary antibodies were used as follows: Rabbit anti-Cux1 (1:250, Santa Cruz, sc-13024), mouse anti-NeuN (1:250, Millipore, MAB377) and DAPI. Appropriate secondary antibodies were used from the Molecular Probes Alexa Series (1:250, Invitrogen). Images were captured by confocal laser scanning microscopy (Zeiss Axio Scope A1, Germany) and image analysis carried out in ImageJ software (111). Areas assessed were found using the following stereotaxic coordinates (110): Bregma anterior-posterior +0.5 mm (frontal motor cortex). Cell counting was performed (Cell Counter, ImageJ) in the frontal motor cortex. For the frontal motor cortex (M1), antibodies Cux1 and NeuN were used to stain pyramidal neurons in layers II-IV; pyramidal neurons in layers V-VI and neurons in general, respectively. Selected images comprised of at least 20 randomly selected microscope fields in the designated layers (area 0.1 mm^2^) from the motor cortex. In addition, DAPI staining allowed the total number of cells to be counted. An average measurement from at least six slices was performed in *Cntn4*^+/+^ (n=5), *Cntn4*^+/-^ (n=4) and *Cntn4*^-/-^ mice (n=4).

### Golgi staining

Golgi staining was performed using a FD Rapid GolgiStain™ kit (FD NeuroTechnologies, Columbia, MD, USA, PK401) according to the instructions of the manufacturer. After treatment with Golgi solutions A, B and C, brains were sectioned with a vibratome (Leica Microsystems, Wetzlar, Germany) coronally at 150 µm thickness from rostral to caudal. The slices were attached to gelatin coated slides and stained with Golgi solutions D + E. Finally, the slices were dehydrated in graded levels of increased concentrations of alcohol, cleaned in xylene, and then cover slipped using Entellan®.

Slices were imaged using light microscopy (Zeiss Axio Scope A2, Germany). Areas assessed were found using the following stereotaxic coordinates (110): Bregma anterior-posterior +0.5 mm (frontal motor cortex). Image analysis was carried out with Golgi Microscope (Zeiss AxioImager M2, Germany) and Neurolucida software (MicroBrightField, Williston, VT, USA). Further slices were imaged using light microscopy (Leica TCS SP8, Leica Microsystems, Mannheim, Germany).

To study the structural differences in spine number and spine morphology caused by *Cntn4*^-/-^ mice, a total of 15 samples of different mice were included for final analysis: *Cntn4*^+/+^, *Cntn4*^+/-^, *Cntn4*^-/-^ mice (N = 5 for each group). A total of 62 pyramidal neurons were included for final analysis (22 for *Cntn4*^+/+^, 16 for *Cntn4*^+/-^, 24 for *Cntn4*^-/-^ neurons). Thin, mushroom, stubby, abnormal and double mushroom spines were counted in the first 25 mm (50 mm to 75 mm) and second 25 mm (the 75 mm to 100 mm) of a branch of the proximal part of the apical dendrite (the first 1/3 part of the apical dendrite) in pyramidal neurons of the CA1 region. The total number of spines includes all morphological categories. Neurolucida and Neuroexplorer software (Plexon, Dallas, Texas, USA), were used for the tracing of spines and analysis (spine number and spine morphology). The different spine morphology categories were counted. Primary branches at the proximal part (the first 1/3 of the apical dendrite) were included for spine analysis and images acquired including circles with a diameter of 200 µm, 150 µm and 100 µm from the branching place. Branches that were not long enough, or with branching places at the branch itself (between 100 and 200 µm) were excluded.

### Neuronal cultures

Neuronal cultures were prepared as described previously (47). P0-P1 mouse cerebral cortices were dissected and dissociated in 0.25% trypsin (PAA) in DMEM/F12 (Gibco, Invitrogen) for 15 min at 37°C. Trypsin was inactivated by adding an equal volume of DMEM/F12 containing 20% FBS (Lonza, Bio Whittaker). Cerebral cortex was dissociated by trituration in DMEM/F12 containing 10% FBS and 20 µg/ml DNase I (Roche) using a fire-polished Pasteur pipette. Dissociated cortical neurons were cultured in Neurobasal medium (Gibco, Invitrogen) on 100 µg/ml poly-L-lysine-coated (Sigma-Aldrich) acid-washed coverslips in a humidified atmosphere with 5% CO_2_ at 37°C.

### Neuronal transfection and analysis

Neurons were transfected as described previously (47). Briefly, at DIV2, neurons in culture were transfected by Lipofectamine LTX according to manufacturer’s protocol (Invitrogen), with a full-length Cntn4 construct or an empty pcDNA3.1 control vector, respectively. At DIV5, neurons were fixed with 4% PFA and 4% Sucrose in PBS, pH 7.4. Immunocytochemistry was performed with the following primary antibodies: rat anti-GFP (Chromotek, 50430-2-AP) 1:500; rabbit anti-Caspase-3 (Cell Signaling Technology, 9664) 1:1000. Images were captured by confocal laser scanning microscopy (Zeiss Axioscop A1). For analysis of neuronal morphological parameters, four independent batch experiments were examined with total analysed cell number n = 279 (*Cntn4* overexpression), n = 439 (empty vector). WIS-Neuromath software (Weizmann Institute) was used for determining morphological parameters (112), which included soma size, number of branches, total dendrite length, and longest dendrite length. For analysis of neuronal apoptosis in the CNTN4 overexpression experiments, the immunoreactivity of caspase-3 in a mean number of 53 transfected neurons was quantified per condition of each independent experiment (n = 4 for empty vector and CNTN4 overexpression, respectively). Positive neurons were analysed by quantification of the number of double-labelled cells as a percentage of the total amount of transfected cells in ImageJ software (111).

### HEK293 cell culture and transfection

HEK293 cells were maintained in high glucose Dulbecco’s modified Eagle’s medium 5 g/L glucose (DMEM; Gibco, UK, 11965084). Cell culture media were supplemented with 10% (v/v) heat-inactivated fetal bovine serum (FBS; Gibco, UK, 11550356), 2 mM L-glutamine (PAA) and 1x penicillin/streptomycin (pen/strep; PAA) and cultured in a humidified atmosphere with 5% CO_2_ at 37°C. HEK293 cells were transfected using polyethylenimine (PEI; Polysciences) (113) or Lipofectamine LTX (Invitrogen, according to manufacturer’s manual).

### Cell adhesion assay

Cell adhesion assays were performed with HEK293 cells as previously described (47,114). HEK293 cells were co-transfected either with pEGFP-N1 (Clontech, 6085-1) or pCAG-DsRed-T1 (gift from Prof. Scheiffele) and full-length pcDNA3.1-FLAG-CNTN4, pSFV-huAPP695 (115) (gift from Prof. De Strooper), pCAG-HA-NLGN1 and pCAG-HA-NRXN1β^-^ (latter two were gifts from Prof. Scheiffele) expression constructs. Separate populations of HEK293 cells were co-transfected either with CNTN4 together with DsRed or with APP together with EGFP expression plasmids. As a positive control, cells were co-transfected with NLGN1 and DsRed or with NRXN1β^-^ and EGFP expression plasmids. NLGN1 and NRXN1β^-^ are well-established *trans*-binding partners (50,51). Since CNTN4 can form homodimers in *trans* (52), cells were also co-transfected with CNTN4 and EGFP or Cntn4 with DsRed expression plasmids, as a second positive control. As a negative control, cells were transfected with either DsRed or EGFP expression plasmids. Cells expressing DsRed, NRXN1β^-^ or APP, were incubated with cells expressing EGFP, NLGN1 or CNTN4. As a negative control, cells expressing DsRed were incubated with cells expressing EGFP. After 48 h, the cells were detached using 1 mM EDTA in PBS, pH 7.4, and centrifuged at 1000 rpm for 5 min. The pellets were resuspended in suspension medium (10% HIFCS, 50 mM Hepes-NaOH pH 7.4, 10 mM CaCl_2_ and 10 mM MgCl_2_ and combined to a total of 5x10^6^ (1:1) in 0.3 ml total volume of 0.5 ml Eppendorf tubes. The cell mixtures were incubated at RT under gentle agitation. The extent of cell aggregation was assessed at 90 min by removing aliquots, spotting them onto culture slides, and imaging by a Leica AF6000 microscope (Leica, UK). The resulting images were then analysed by counting the number and size of particles using ImageJ. An arbitrary value for particle size was then set as a threshold based on negative control values. The aggregation index was calculated by expressing the number of particles participating in aggregation as a percentage of the total particles in 5-10 fields of 1.509 mm^2^ per cell suspension combination of each independent experiment (n=3).

### Cell surface binding assay

To investigate whether CNTN4 interacts with APP, a cell surface binding assay was used with slight modifications (116). Transfection of HEK293 cells with pIGplus-RGMa-Fc or pSFV-huAPP695 was performed. Forty-eight hours after transfection the medium with soluble RGMA-Fc or APP was concentrated through a 50,000 kDa column (YM-50, Millipore). The concentrated proteins were supplemented with Dulbecco’s modified Eagle’s medium 1 g/L glucose (DMEM; Gibco, Invitrogen) with 2 mM L-glutamine (PAA) and 1x penicillin/streptomycin (pen/strep; PAA) and distributed in 6-well plates with HEK293 cells transfected with Neo1.a-AP-His (Addgene #71963) or pcDNA3.1-CNTN4-FLAG. Binding between the proteins was allowed overnight in a humidified atmosphere with 5% CO_2_ at 37°C. Cells were fixed with 4% PFA in PBS, pH 7.4, and 0.01% sodium azide. Immunocytochemistry was performed with the following primary antibodies: rabbit anti-FLAG (20543-1-AP, Proteintech) 1:200; rabbit anti-His (10001-0-AP, Proteintech) 1:200, goat anti-APP (AF1168, R&D Systems) 200 μg/mL, mouse anti-Fc (M4280, Sigma Aldrich) 1:100. Appropriate secondary antibodies were used from the Molecular Probes Alexa Series (1:250, Invitrogen). For the cell surface binding analysis, images from the Leica AF6000 microscope (Leica, UK) were used. Analyses were performed of about 300 transfected cells per condition of each independent experiment (n=3). The images were analysed by quantification of the number of double labelled cells as a percentage of the total amount of transfected cells in ImageJ. Statistical analysis was carried out using unpaired Student’s *t*.

### Immunoprecipitation

Immunoprecipitation (IP) experiments were performed using GFP-Trap-A beads (Chromotek, gta), according to manufacturer’s manual, as previously described (47). A biotin-and GFP-tagged extracellular rat *Cntn4* (CNTN4-TMGFPBio) fusion protein was generated by subcloning the coding sequence of the extracellular CNTN4 domains (NM_053879.1: nt 476-3394), excluding the coding sequence of the GPI anchor. This was amplified from wild-type *Cntn4* cDNA (BluescriptIISK-CNTN4) and ligated to the sequence of plexin-A1 transmembrane domain coding sequence (NM_008881.2: nt 3962-4123). The coding sequences of a five glycine linker and intracellular GFP and biotin tags followed and were inserted in a pcDNA3.1(-)/myc-His (Invitrogen) vector backbone. The control vector (TMGFPBio) is identical, but it is truncated beyond the transmembrane domain.

For proteomics, HEK293 cells expressing the indicated GFP-tagged fusion proteins were collected in ice-cold PBS and centrifuged at 1000 rpm in a precooled centrifuge at 4°C for 5 min. Cell pellets were lysed in lysis buffer [10 mM Tris-HCl, pH 7.5, 150 mM NaCl, 0.5 mM EDTA, 0.5% NP40, 1 mM PMSF and Complete protease inhibitor cocktail (Roche)], incubated on ice for 30 min and centrifuged at 13,200 rpm at 4°C for 10 min. Cleared supernatant containing roughly 5.4-6.6 mg of protein was mixed with 50 µl GFP-Trap-A agarose beads (Chromotek), which had been equilibrated in dilution buffer [10 mM Tris-HCl, pH 7.5, 150 mM NaCl, 0.5 mM EDTA, 1 mM PMSF and Complete protease inhibitor cocktail (Roche) at 4°C. After 1.5 h incubation at 4°C, beads were washed two times in dilution buffer. Precipitated proteins were eluted by boiling the pull-down samples in NuPAGE LDS sample buffer (Invitrogen) containing 2% β-mercaptoethanol at 95°C for 10 min.

Thirty microliter of each sample was ran on a 12% Bis-Tris 1D SDS-PAGE gel (Biorad) either for 2-3 cm or ran completely and stained with colloidal coomassie dye G-250 (Gel Code Blue Stain Reagent, Thermo Scientific). The lane was cut into bands, which were treated with 6.5 mM dithiothreitol (DTT) for 1 h at 60°C for reduction and 54 mM iodoacetamide for 30 min for alkylation. The proteins were digested overnight with trypsin (Promega) at 37°C. The peptides were extracted with acetonitrile (ACN) and dried in a vacuum concentrator.

### Mass Spectrometry: RP-NanoLC-MS/MS

Mass spectrometry data was acquired as previously described (47). Data was acquired using an LTQ-Orbitrap coupled to an Agilent 1200 system or an Obritrap Q Exactive mass spectrometer connected to an Agilent 1290 system. In case of the LTQ-Orbitrap, peptides were first trapped ((Dr Maisch GmbH) Reprosil C18, 3 µm, 2 cm x 100 µm) before being separated on an analytical column (50 µm x 400 mm, 3 µm, 120 Å Reprosil C18-AQ). Trapping was performed at 5 µl/min for 10 min in solvent A (0.1 M acetic acid in water) and the gradient was as follows; 10-37% solvent B (0.1 M acetic acid in 80% acetonitrile) in 30 min, 37-100% B in 2 min, 100% B for 3 min, and finally solvent A for 15 min. The flow was passively split to 100 nl min^-1^. Data was acquired in a data-dependent manner, to automatically switch between MS and MS/MS. Full scan MS spectra from *m/z* 350 to 1500 were acquired in the Orbitrap at a target value of 5e5 with a resolution of 60,000 at *m/z* 400 in case of the LTQ-Orbitrap XL and 30,000 for the LTQ-Discovery. The five most intense ions were selected for fragmentation in the linear ion trap at a normalized collision energy of 35% after the accumulation of a target value of 10,000. In case of the Q Exactive samples were first trapped ((Dr Maisch GmbH) Reprosil C18, 3 µm, 2 cm x 100 µm) before being separated on an analytical column (Agilent Poroshell EC-C18, 2.7 µm, 40 cm x 50 µm). Trapping was performed for 10 min in solvent A and the gradient was as follows: 13-41% solvent B in 35 min, 41-100% in 3 min and finally solvent A for 10 min. Flow was passively split to 100 nl min^-1^. The mass spectrometer was operated in data-dependent mode. Full scan MS spectra from *m/z* 350-1500 were acquired at a resolution of 35,000 at *m/z* 400 after accumulation to a target value of 3e6. Up to ten most intense precursor ions were selected for fragmentation. HCD fragmentation was performed at normalised collision energy of 25% after the accumulation to a target value of 5e4. MS/MS was acquired at a resolution of 17.500. In all cases nano-electrospray was performed at 1.7 kV using an in-house made gold-coated fused silica capillary (o.d. 360 µm; i.d. 20 µm; tip i.d. 10 µm).

### Proteomics data analysis

Raw files were processed using Proteome Discoverer 1.3 (version 1.3.0.339, Thermo Scientific Bremen, Germany). The database search was performed against the Swissprot database (version August 2014) using Mascot (version 2.4.1, Matrix Science, UK) as search engine. Carbamidomethylation of cysteines was set as a fixed modification and oxidation of methionines was set up as a variable modification. Trypsin was specified as enzyme and up to two miss cleavages were allowed. Data filtering was performed using percolator, resulting in 1% false discovery rate (FDR). An additional filter was Mascot ion score >20. Raw files corresponding to one sample were merged into one result file. Data was further analysed with Saint (48) using the Crapome web interface in order to identify interacting proteins. Default settings were used for calculating the FC-A and FC-B score. The probability score was calculated using Saint Express performing 20,000 iterations.

### SH-SY5Y cell culture and CRISPR/Cas9 mediated knock out (KO) of *CNTN4* and ***APP***

Neuroblastoma SH-SY5Y cells were cultured in Dulbecco’s modified Eagle’s medium with GlutaMAX (DMEM/F-12, Gibco, UK, 10566016) supplemented with 10% (v/v) heat-inactivated fetal bovine serum. CRISPR/Cas9 gene editing was performed as described elsewhere (117). In brief, sgRNAs were designed in Benchling (Benchling Software, 2020), ordered from Integrated DNA Technologies and cloned into pSpCas9(BB)-2A-GFP(PX458) and pU6-(BbsI)_CBh-Cas9-T2A-mCherry. pSpCas9(BB)-2A-GFP (PX458) was a gift from Feng Zhang (Addgene plasmid #48138; http://n2t.net/addgene:48138; RRID:Addgene_48138). pU6-(BbsI)_CBh-Cas9-T2A-mCherry was a gift from Ralf Kuehn (Addgene plasmid #64324; http://n2t.net/addgene:64324; RRID:Addgene_64324). Paired sgRNAs targeting regions of *CNTN4* and *APP* were designed to generate homozygous knockout in addition to the non-targeting empty vector control (full sequence detail in Table S1, Supplementary Materials). sgRNA plasmids validated by Sanger sequencing were transfected into SH-SY5Y cells using nucleofection (SF Cell Line 4D-Nucelofector X kit, Lonza, Germany, V4XC-2012) according to the manufacturer’s protocol. After low-density seeding, single clones were isolated and expanded in 96-well plates. Overall, we sequenced at least ten clones per construct and confirmed homozygous single cell clones at least twice by Sanger sequencing. Both indels induced premature stop codons as validated by PCR and Sanger sequencing. *CNTN4* and *APP* deficiency was further shown on RNA and protein level using real-time RT-PCR and Western blot, respectively. RNA extraction, cDNA generation, and real-time RT-PCR as well as protein extraction and Western blots were performed as previously described (44).

### *In vitro* cell morphology assay

SH-SY5Y cells were differentiated using a continuous application of retinoic acid (RA, Sigma Aldrich, UK) and brain-derived neurotrophic factor (BDNF, Peprotech, UK, 450-02). Differentiation media consisted of DMEM/F-12 (0% FBS), 10 µM RA and 20 ng/mL BDNF. Cells were differentiated for five days changing the medium every other day. On day 5, cells were fixed with 4% PFA in PBS, pH 7.4. Cells were permeabilised and blocked for 1 hour at room temperature. Immunocytochemistry was performed with primary antibody mouse anti-β(III)-Tubulin (R&D Systems, MAB1195) 1:500, with appropriate secondary antibody from the Molecular Probes Alexa Series (1:250, Invitrogen). Nuclei were counter-stained with DAPI. Images were captured by fluorescence microscope (Leica DM4000 LED) with LAS-X software. For analysis of neuronal morphological parameters, three independent batch experiments were examined with total analysed cell number at least n = 90 per genotype. Image analysis was carried out in ImageJ, which included cell area, total and longest neurite length.

### Statistical analysis

For statistical analysis, data was plotted as the mean ± standard error of the mean, unless otherwise stated. Statistical analysis was carried out using ANOVA and unpaired Student’s *t* tests where appropriate (Graphpad Prism). Further details are provided in the Supplementary Materials.

## Supporting information

Supplemental Datasheet

## Acknowledgements

We thank Dr. Yoshihiro Yoshihara, RIKEN BSI, for providing the *Cntn4* mice and helpful advice. We thank Mr. Henk Spierenburg for performing the genotyping of the animals. We thank Canon foundation in Europe, ARUK South West Network, QUEX Initiator, British Society for Neuroendocrinology, and Northcott Devon Medical Foundation for financial support.

