## Supplementary figures and images for "CNTN4 modulates neural elongation through interplay with APP"

### Supplemental Figure 1

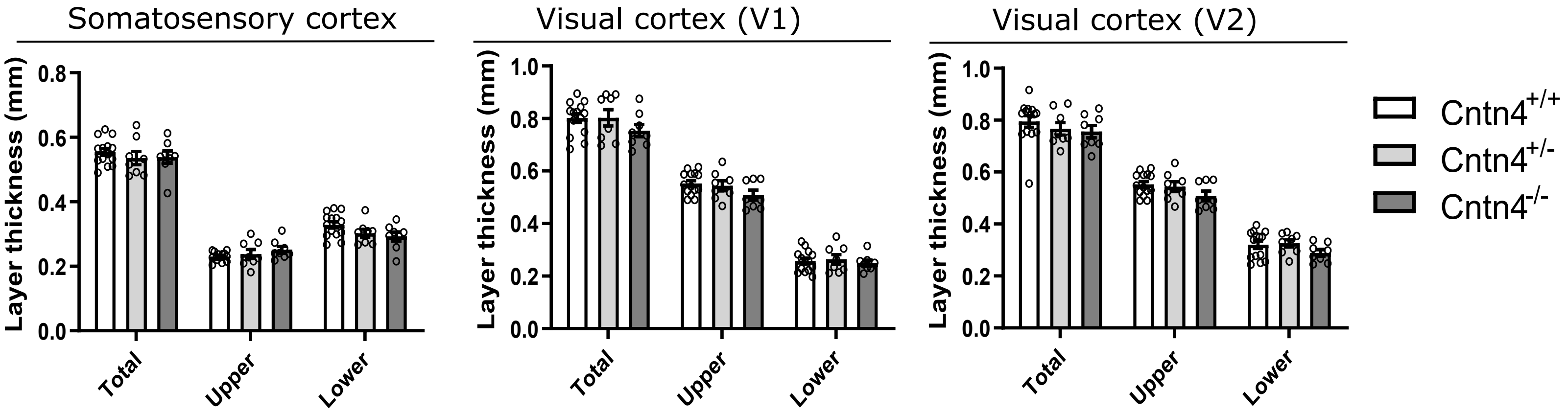

### Supplemental Figure 2

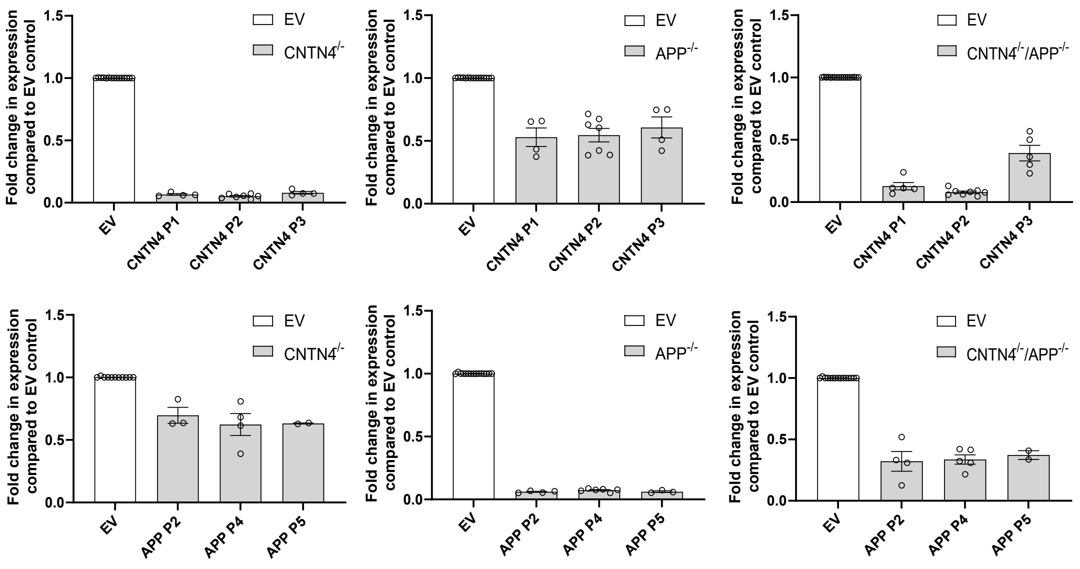

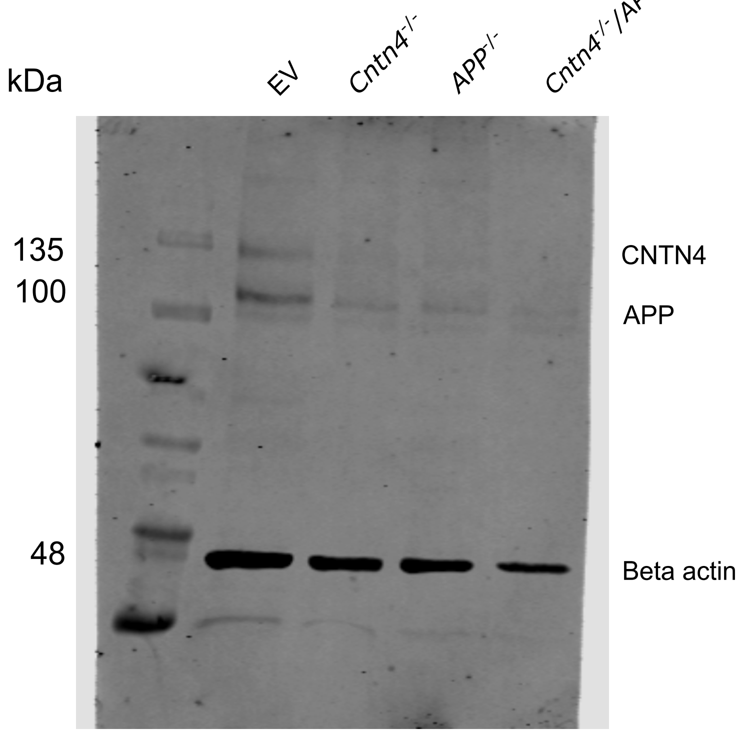
