## Supplemental Information for "CNTN4 modulates neural elongation through interplay with APP"

**Supplementary Information**

**Materials and Methods**

**Animals**

*Cntn4*-deficient mice were kindly provided by Dr. Yoshihiro Yoshihara (RIKEN, Japan) (45). These mice were generated using a standard gene-targeting method as previously described. A targeting vector was designated to mutate the translation start codon (ATG) in the exon 2 of the Cntn4 gene into a stop codon (TAG) and to introduce a pgk-neo selection marker. Consequently, these mice were backcrossed with C57BL/6 mice more than nine times. Upon arrival in the University Medical Center Utrecht (UMCU, Utrecht), the mice were re-derived, followed by heterozygous breeding to generate littermate wild types (*Cntn4*^+/+^), heterozygotes (*Cntn4*^+/-^) and homozygous *Cntn4* knockout mice (*Cntn4*^-/-^) for the use in our experiments.

Genotyping was carried out on 6-week old mice. Briefly, DNA from ear tag was extracted following lysis (0.01 M NaOH, 1 mM EDTA, Proteinase K), and subjected to PCR using specific primers for *Cntn4* (Table S1) with the resulting products separated and visualized using agarose gel electrophoresis. Genotype was determined by the presence of product molecular weight 1100 bp (*Cntn4^+/+^*), 850 bp (*Cntn4^-/-^*) or the presence of both products (*Cntn4^+/-^*).

All mice were kept on a normal day/night cycle and had access to food and water *ad libitum* (UMCU, Utrecht). For immunohistochemistry, adult male mice were anesthetized with an overdose of sodium pentobarbital (19.4 µl/gr) and were perfused intracardially with 0.9% saline, followed by 4% PFA in phosphate-buffered saline (PBS), pH 7.5. Brains were post fixed in 4% PFA before transferring to 30% sucrose for cryopreservation.

**PCR and Sanger sequencing**

SH-SY5Y clonal cells were screened via PCR using specific primers for *CNTN4* and *APP* (Table S1) with the resulting products separated and visualized using agarose gel electrophoresis. *CNTN4* genotype was determined by the presence of product molecular weight 341 bp (*CNTN4^+/+^*), 249 bp (*CNTN4^-/-^*) or the presence of both products (*CNTN4^+/-^*). *APP* genotype was determined by the presence of product molecular weight 2905 bp (*APP ^+/+^*), 701 bp (*APP ^-/-^*) or the presence of both products (*APP ^+/-^*). Sanger sequencing of PCR amplicon confirmed indel (Genewiz, UK).

**Quantitative real-time PCR assays of mRNA expression**

The mRNA expression of *Cntn4* in cortex extracted from adult male mice (Zhang et al., 2014) was measured by real-time PCR (RT-PCR), as previously reported (Oguro-Ando et al., 2021). The mRNA expression of *CNTN4* and *APP* in SH-SY5Y cells (Zhang et al., 2014) was measured by real-time PCR (RT-PCR) using PrimeScript reverse transcription reagent kit (Takara Bio Group, Japan, RR037A), as described (Oguro-Ando et al., 2015). Briefly, total RNA was extracted from SH-SY5Y cells using Trizol reagent (Invitrogen Corp., CA, USA, R2050-1-50) and Direct-zol RNA miniprep (Zymo Research, CA, USA, R2051) according to manufacturer’s instructions. Total RNA was converted to cDNA using PrimeScript, and RT-PCR was performed using HOT FIREPol EvaGreen (Solis Biodyne, Estonia, 08-24-00001) on the QuantStudio 6 Flex real-time PCR system (ThermoFisher Scientific, UK). *GAPDH and POLR2A* mRNAs were employed as endogenous controls. The primers employed in these RT-PCR experiments are described in Table S1. The relative expression of each test transcript was determined by the comparative Ct approach. Expression levels were calculated relative to the geometric mean of the endogenous controls, but also to the global mean of expression across all transcripts tested, which was empirically determined to vary across test conditions to provide a robust baseline. Expression levels were normalized to the mean of expression seen in empty vector control SH-SY5Y cells. Analysis was performed on three SH-SY5Y cell lines (*CNTN4^-/-^, APP^-/-^, CNTN4^-/-^/APP^-/-^*). Statistical analysis between SH-SY5Y cell lines was performed using one-way ANOVA and Tukey’s multiple comparison post-hoc test. Data are expressed as means ± S.E.M.

**Western blotting**

The protein expression of CNTN4 in cortex extracted from adult male mice was measured by Western blotting as previously reported (Oguro-Ando et al., 2021). SH-SY5Y cells were re-suspended in ice-cold RIPA lysis buffer (10 mM Tris/HCl pH 7.5, 150 mM NaCl, 0.1% SDS, 1% Triton X-100, 1% Deoxycholate, 0.5 mM EDTA, 1 mM PMSF, Complete protease inhibitor cocktail (Roche, UK), protease inhibitor cocktail (Sigma, UK) and phosphatase inhibitor cocktail 2, 3 (Sigma, UK), placed on ice for 10 mins, followed by centrifugation at 13,200 rpm for 10 min at 4°C. The supernatant was collected, SDS sample buffer containing 2% β-mercaptoethanol was added and samples were boiled for 5 min at 95°C. Proteins were separated in 7.5% SDS-PAGE gels and transferred onto PVDF membrane (Immobilon-P, Merck Millipore, UK). Membranes were incubated in blocking buffer (TBS, 1% (v/v) Tween 20, 5% milk powder, 5% BSA, 1% FBS) for 1 hour at RT. Membranes were incubated with corresponding primary antibodies in blocking buffer (TBS, 1% (v/v) Tween 20, 2% milk powder) overnight at 4°C. Primary antibodies used: goat anti-CNTN4 (EB11768, Everest, UK) 1:1000, rabbit anti-APP (AF1168, Abcam, UK) 1:1000 and mouse anti-beta-actin (NB600-501, Novus Biologicals, UK) 1:2500. Secondary antibodies used: donkey anti-goat (IRDye 800CW, LI-COR Biosciences, NE, USA) 1:5000 and goat anti-mouse (DyLight 680, Invitrogen, UK) 1:5000. Blots were imaged using the LI-COR Odyssey CLx (LI-COR Biosciences, NE, USA). Western blotting was performed on four SH-SY5Y cell lines (empty vector, *CNTN4^-/-^, APP^-/-^, CNTN4^-/-^/APP^-/-^*).

**Statistical analysis**

For cortical layer thickness, measurements were performed, in each area, on at least three slices in *Cntn4*^+/+^ (n=7), *Cntn4*^+/-^ (n=4) and *Cntn4*^-/-^ mice (n=4). Statistical analyses were carried out using one-way ANOVA and Tukey’s multiple comparison post-hoc test.

For immunohistochemistry, measurements from at least six slices was performed in *Cntn4*^+/+^ (n=5), *Cntn4*^+/-^ (n=4) and *Cntn4*^-/-^ mice (n=4). Statistical analyses were carried out using one-way ANOVA and Tukey’s multiple comparison post-hoc test.

For *Cntn4* Golgi analysis, Apical: Neurite length (*Cntn4*^+/+^ 23, *Cntn4*^+/-^ 22, *Cntn4*^-/-^ 21); DCI (*Cntn4*^+/+^ 22, *Cntn4*^+/-^ 22, *Cntn4*^-/-^ 23); Tips (*Cntn4*^+/+^ 22, *Cntn4*^+/-^ 22, *Cntn4*^-/-^ 23); Total intersection (*Cntn4*^+/+^ 23, *Cntn4*^+/-^ 22, *Cntn4*^-/-^ 23); Last Sholl intersection (*Cntn4*^+/+^ 23, *Cntn4*^+/-^ 22, *Cntn4*^-/-^ 23). Basal: Neurite length (*Cntn4*^+/+^ 23, *Cntn4*^+/-^ 24, *Cntn4*^-/-^ 22); DCI (*Cntn4*^+/+^ 22, *Cntn4*^+/-^ 24, *Cntn4*^-/-^ 22); Tips (*Cntn4*^+/+^ 22, *Cntn4*^+/-^ 24, *Cntn4*^-/-^ 22); Total intersection (*Cntn4*^+/+^ 23, *Cntn4*^+/-^ 23, *Cntn4*^-/-^ 23); Last Sholl intersection (*Cntn4*^+/+^ 23, *Cntn4*^+/-^ 23, *Cntn4*^-/-^ 23) Spines: 5 mice per genotype (*Cntn4*^+/+^ 22, *Cntn4*^+/-^ 16, *Cntn4*^-/-^ 24). Statistical analyses were carried out using one-way ANOVA and Tukey’s multiple comparison post-hoc test.

For *APP* Golgi analysis: 5 mice per genotype (APP^+/+^ 30, APP^-/-^ 30). Statistical analyses were carried out using unpaired Student’s *t* test.

Neuronal transfection analysis: 4 batches (155, 78, 138, 68 empty vector cells) and (92, 23, 99, 65 CNTN4 overexpressing cells). Average of 90 cells analysed per condition. Total analysed cell number n = 439 (CNTN4 OE), n = 279 (empty vector). Statistical analyses were carried out using unpaired Student’s *t* test.

Cell adhesion assay: Total analysed image frames per condition, n = 42 (dsRed-EGFP), n = 63 (CNTN4-APP), n = 43 (CNTN4-CNTN4), n = 41 (NRXN-NLGN); across 3 independent experiments. Statistical analyses were carried out using one-way ANOVA and Tukey’s multiple comparison post-hoc test.

Table 1: Primer sequences

|  | Gene | Forward Primer (5’ to 3’) | Reverse Primer (5’ to 3’) |
| --- | --- | --- | --- |
| *i)* | *Cntn4* | TGG TAG ATG GAT CGA TGG CAA ACA TG  AGC CCC AGT TTT TGC CTA AGC AT | TTC ATC ACT CCT GAA TCA CAC ATG TCA |
| *ii)* | *CNTN4* | GGCGATTATCCTGATAGGAA | CACGGTATTGGTAACCACAC |
|  | *APP* | GCACTTGTCAGGAACGAGAA | GCGGAATTGACAAGTTCCGAG |
| *iii)* | *CNTN4* | ACAGTAGAGTTCCAGTGCCA | AGCTTCATGTGCCAACAACA |
|  | *APP* | AGGGACAATTTCACTTGCAGC | TGACCAAAACAGAACGCCAT |
| *iv)* | *CNTN4* primer set 1 | GGGCCACCTACACCACTAAT | CATTCCAGCTTCACCGTTGC |
|  | *CNTN4* primer set 2 | GGTAACGCATGATCACTCGC | ATCACCAGCTGAATCCTGCC |
|  | *CNTN4* primer set 3 | Sigma KiCqStart® SYBR®Green Primers. HA14242300, FH2_CNTN4, Species (human), Primer Pair ID 1: H_CNTN4_2 | Sigma KiCqStart® SYBR®Green Primers. HA14242301, RH2_CNTN4, Species (human), Primer Pair ID 1: H_CNTN4_2 |
|  | *APP* primer set 1 | GCCAACCAACCAGTGACCAT | CTCACCAACTAAGCAGCGGT |
|  | *APP* primer set 2 | GCTGGTGGAGACACACATGGCC | GGATCTGAGCGGCTTTCTTGGG |
|  | *APP* primer set 3 | Sigma KiCqStart® SYBR®Green Primers. HA14242298, FH1_APP, Species (human), Primer Pair ID 1: H_APP_1 | Sigma KiCqStart® SYBR®Green Primers. HA14242299, RH1_APP, Species (human), Primer Pair ID 1: H_APP_1 |
|  | *GADPH* | TCCTCTGACTTCAACAGCGAC | GCTGTAGCCAAATTCGTTGTCA |
|  | *POLR2A* | CCATCAAGAGAGTCCAGTTCG | ACCCTCCGTCACAGACATTC |

Primer sequences used for *i)* determining *Cntn4* genotype in mice. Paired sgRNA sequences ii) for knockout of *CNTN4* and *APP*, and primer sequences used for iii) determining genotype in human SH-SY5Y cells. Primer sequences used for iv) quantifying *CNTN4* and *APP* mRNA expression in human SH-SY5Y cells using the PrimeScript reverse transcription kit (Takara Bio Group).

**Figure S1:** Equal cortical thickness between Nissl-stained sections of adult *Cntn4^+/+^*, *Cntn4^+/-^* and *Cntn4^-/-^* mice. (A) Quantitative analysis of the somatosensory and visual cortices layer thickness (upper I-IV, lower V-VI and total) demonstrate no significant differences in thickness in all layers between *Cntn4^+/+^* and *Cntn4^-/-^* mice. Analysis was performed, in each area, on at least two slices in *Cntn4^+/+^* (n=7), *Cntn4^+/-^* (n=4) and *Cntn4^-/-^* mice (n=4). Data are presented as mean ± S. E. M, p > 0.05 one-way ANOVA.

**Figure S2**: Expression levels in CNTN4 and APP deficient cells A) Fold change of *Cntn4* and *APP* mRNA expression levels in *CNTN4^-/-^, APP^-/-^, CNTN4^-/-^/APP^-/-^* cells compared to empty vector cells, generated by qRT-PCR. Analysis was performed on three SH-SY5Y cell lines (*CNTN4^-/-^, APP^-/-^, CNTN4^-/-^/APP^-/-^*) using one-way ANOVA and Tukey’s multiple comparison post-hoc test. B) Protein extracted from SH-SY5Y cells was analyzed by Western blot. Blots stained with anti-CNTN4 and anti-APP antibodies revealed lower expression in *CNTN4^-/-^, APP^-/-^, CNTN4^-/-^/APP^-/-^* cells, respectively. Western blotting was performed on four SH-SY5Y cell lines (empty vector, *CNTN4^-/-^, APP^-/-^, CNTN4^-/-^/APP^-/-^*). Molecular weights are as follows: CNTN4 = 150 kDa; APP = 100-140 kDa; beta-actin = 47 kDa.
